# What influences selection of native phosphorelay architectures?

**DOI:** 10.1101/2020.05.21.108001

**Authors:** Rui Alves, Baldiri Salvado, Ron Milo, Ester Vilaprinyo, Albert Sorribas

## Abstract

Phosphorelays are signal transduction circuits that combine four different phosphorylatable protein domains for sensing environmental changes and use that information to adjust cellular metabolism to the new conditions in the milieu. Five alternative circuit architectures account for more than 99% of all phosphorelay operons annotated in over 9000 fully sequenced genomes, with one of those architectures accounting for more than 72% of all cases.

Here we asked if there are biological design principles that explain the selection of preferred phosphorelay architectures in nature and what might those principles be. We created several types of data-driven mathematical models for the alternative phosphorelay architectures, exploring the dynamic behavior of the circuits in concentration and parameter space, both analytically and through over 10^8^ numerical simulations. We compared the behavior of architectures with respect to signal amplification, speed and robustness of the response, noise in the response, and transmission of environmental information to the cell.

Clustering analysis of massive Monte Carlo simulations suggests that either information transmission or metabolic cost could be important in selecting the architecture of the phosphorelay. A more detailed study using models of kinetically well characterized phosphorelays (Spo0 of *Bacillus subtilis* and Sln1-Ypd1-Ssk1-Skn7 of *Saccharomyces cerevisiae*) shows that information transmission is maximized by the natural architecture of the phosphorelay. In view of this we analyze seventeen additional phosphorelays, for which protein abundance is available but kinetic parameters are not. The architectures of 16 of these are also consistent with maximization of information transmission.

Our results highlight the complexity of the genotype (architecture, parameter values, and protein abundance) to phenotype (physiological output of the circuit) mapping in phosphorelays. The results also suggest that maximizing information transmission through the circuit is important in the selection of natural circuit genotypes.

## Background

Organisms and cells use signal transduction circuits to detect environmental changes and make decisions on how to adjust their internal milieu to better survive those changes. Those circuits modulate the cellular response and its metabolic adjustments.

Phosphorelays (PR) are a large and important class of signal transduction circuits in microorganisms and some plants^1–4^. A self-phosphorylating Sensor Kinase (SK) modulates its own phosphorylation state in response to the environmental signal. The phosphorylated SK transfers its phosphate to an aspartate residue in a Response Regulator (RR) protein. This response regulator transfers its phosphate to a Histidine phosphotransfer (Hpt) protein domain, which then transfers the phosphate to a final RR (Figure 1). The phosphorylation state of the circuit modulates cellular activity and adaptation. PR are important for making life or death decisions about sporulation ^5^, in adapting to various stressors ^6^, such as changing levels of oxygen ^7^, or in developmental decisions made by many plants ^8^.

**Figure 1.**
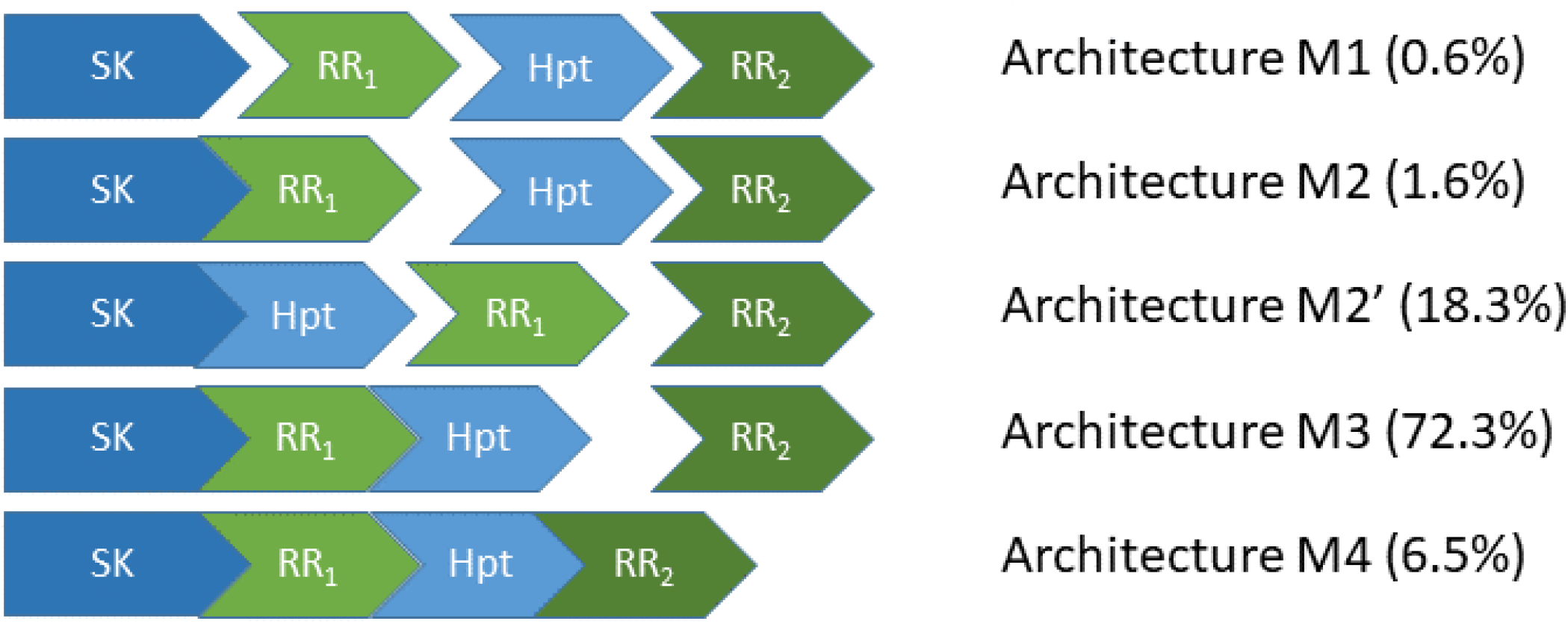
The five most abundant phosphorelay circuit architectures, as inferred from operon structure, account for over 99% of all detected phosphorelays. Architecture M1 is for a circuit where the four phosphorylatable domains exist in independent proteins. Architecture M2 is for a circuit with a hybrid Sensor Kinase (SK), which contains the SK and the first Response Regulator (RR_1_) domain in the same protein, while the remaining phosphorylatable domains exist in independent proteins. Architecture M2’ is for a circuit where the SK and the Hpt domains are in the same protein, while both RR domains exist in independent proteins. Architecture M3 is for a circuit where the SK, RR_1_ and the Hpt domains are in the same protein, while the final RR, RR_2_, is in an independent protein. Architecture M4 is for a circuit where all phosphorylatable domains exist in the same protein. A total of 5219 PR operons were surveyed, out of which 5182 fall in one of the five architectures shown here.

The four-step nature of the PR phosphotransfer cascade creates steeper signal response curves than those of two component systems (TCS), a simpler signal transduction alternative to PR with only two phosphotransfer steps ^9–15^. This difference could explain the evolution of PR as an alternative to TCS signaling.

Still, PR circuits can assemble the four-step phosphotransfer cascade using alternative protein domain assembly architectures ^12–14,16–18^. This raises the question of whether alternative PR architectures may have irreducible physiological differences for the behavior of the circuit or if those differences may be quenched easily by evolving appropriate parameter values ^9,10,19–21^. In this context, can we identify general physiological requirements that dominate the selection of PR circuit architectures and/or parameters?

To address this question we used a comprehensive census of PR circuit architectures, and analyzed over 9000 fully sequenced microbial and plant genomes. We found that more than 72% of all PR coded in a single operon have the same circuit architecture (M3 of Figure 1) ^16^. Four additional architectures account for another 27% of all other PR (architectures M1, M2, M2’, an M4 of Figure 1) ^16^.

To understand the physiological differences between the five PR architectures we consider a set of physiological variables that are important for the function of PR. We analyze how adjusting those variables for improved circuit performance depends on the architecture, the protein abundance, and the parameter values of the PR. Our results suggest that optimizing different aspects of environmental information transmission to the cell could be crucial for selection of a specific PR architecture. They also suggest that fine tuning of parameter values and protein abundances could, in many cases, overcome intrinsic differences between the physiological responses of alternative circuit architectures. Together, the results reported here have consequences for understanding the evolution of PR signal transduction circuits and for the development of synthetic biology PR circuit applications.

## Results

### Physiological variables as a proxy for circuit performance in signal transduction

The overall fitness of organisms is affected by how their molecular components organize into biological circuits ^22,23^. The organization of molecular circuits with common biological functions in different organisms may have alternative architectures. Often, these alternatives lead to improved circuit performance in the context of the organisms where they are observed. Understanding how the alternative architectures affect circuit performance requires that the physiological variables that are important for the biological function of the circuit are understood. Over the last few decades, several of these physiological variables were identified as important in determining the performance of molecular signal transduction circuits. We compiled those variable and summarize them in Table 1.

**Table 1.** Physiological variables used as proxy for performance in signal transduction circuit.

| Variables | Circuit performance improves with*: | Experimental support |
| --- | --- | --- |
| Signal amplification | Higher amplification | 9,10,36,69–72 |
| Noise attenuation<sup>^</sup> | Attenuated noise | 57,72–74 |
| Information transmission<sup>^</sup> | Higher transmission | 57,72–74 |
| Robustness to changes in parameter values | High robustness (low sensitivity) | 9,10,36,58,69 |
| Speed of response to changes | Rate of adaptation | 9,10,29,36,48,71,75 |
| Metabolic cost of circuit | Low cost | 76,77 |

\*as a general trend; <sup>^</sup> Relevant in the stochastic domains of dynamic behavior.

### Comparing physiological variables between architectures

To investigate if any physiological variable or combination thereof might explain the genomic frequency differences between the five alternative PR architectures shown in Figure 1, we started by creating mathematical models that permit estimating the variables from Table 1 for each of the architectures. Then, we analyzed those models and compared the variables between architectures in a controlled manner.

We note that the comparisons consider variables that are physiologically relevant in two domains for the dynamic response of the circuits (Table 1): (i) the deterministic domain (≥~1000 copies of the circuit per cell); (ii) the stochastic domain (<1000 copies of the circuit per cell). The methods used to calculate the different physiological variables depend on the regime being analyzed.

In the deterministic domain, signal amplification, robustness, speed, and metabolic cost can be calculated analytically for each architecture (see Datafile_S1.zip for the analytical expressions). We compare which architecture has the highest value for each of these variables by taking the ratio of that variable between all possible pairs of architectures. A detailed analysis of the analytical ratios shows that the best performing architecture with respect to any variable depends on the parameter values.

To investigate if, statistically, there were physiological differences between architecture with respect to the different physiological variables we used a Monte Carlo approach. We randomly generated sets of parameter values, tested the system they created and selected those that create realistic behavior for each architecture and permit comparing that behavior between architectures in a controlled manner (see methods for details). We stopped generating new sets when five thousand realistic systems were obtained for each architecture. Overall, we scanned for more than one hundred and sixty million parameter sets.

Then, for each variable and set of parameter values, we calculated the numerical value of the physiological variables. Finally, for each variable, we selected the 10% of systems that performed better and calculated the percentage of each architecture contained in this high-performance set.

The percentages of systems for each architecture in the best performing systems are very different from the relative genomic frequency of the architectures. Because of this we ranked the five architectures from best (1) to worst (5). Results are summarized in Table 2.

**Table 2.** Comparative results for the various physiological variables. Dark green indicate best performance. Dark yellow indicates worst performance.

| Physiological Variable | M1 | M2 | M2' | M3 | M4 |
| --- | --- | --- | --- | --- | --- |
| Signal amplification | 4.3 % | 10.3 % | 36.2 % | 33.8 % | 15.4 % |
| Robustness | 21.6 % | 26.1 % | 18.5 % | 17.5 % | 16.3 % |
| Intrinsic Noise SK | 1.6 % | 5.4 % | 8.6 % | 30.3 % | 54. % |
| Intrinsic Noise RR2 | 19.8 % | 17.8 % | 18.1 % | 23.5 % | 20.8 % |
| INRR2/INSK | 6.1 % | 15.3 % | 15.1 % | 31.2 % | 32.3 % |
| SK->RR1 MI | 28.9 % | 24.9 % | 24 % | 13.3 % | 9 % |
| SK->RR2 MI | 29.4 % | 24.3 % | 24.5 % | 11.1 % | 10.7 % |
| SK->RR2/SK->RR1 MI | 9.6 % | 8.8 % | 18.5 % | 14.9 % | 48.2 % |
| Response time | 29. % | 21.9 % | 25.4 % | 14. % | 9.6 % |
| Cost | 1.9 % | 5.1 % | 8.3 % | 31.2 % | 53.4 % |
| **Genomic frequency** | **0.6 %** | **1.6 %** | **18.3 %** | **72.3 %** | **6.5 %** |

| Physiological Variable | Rank M1 | Rank M2 | Rank M2' | Rank M3 | Rank M4 |
| --- | --- | --- | --- | --- | --- |
| Signal amplification | 5 | 4 | 1 | 2 | 3 |
| Robustness | 2 | 1 | 3 | 4 | 5 |
| Intrinsic Noise SK | 5 | 4 | 3 | 2 | 1 |
| Intrinsic Noise RR2 | 3 | 5 | 4 | 1 | 2 |
| INRR2/INSK | 5 | 3 | 4 | 2 | 1 |
| SK->RR1 MI | 1 | 2 | 3 | 4 | 5 |
| SK->RR2 MI | 1 | 2 | 2 | 4 | 5 |
| SK->RR2/SK->RR1 MI | 4 | 5 | 2 | 3 | 1 |
| Response time | 1 | 3 | 2 | 4 | 5 |
| Cost | 5 | 4 | 3 | 2 | 1 |
| **Genomic frequency rank** | **5** | **4** | **2** | **1** | **3** |

We then clustered the vectors of architecture ranks (Figure 2) using a Canberra distance metric. We find that Information transmission and cost are the physiological variable that generate a rank that is the most similar to that generated by genomic frequency.

**Figure 2.**
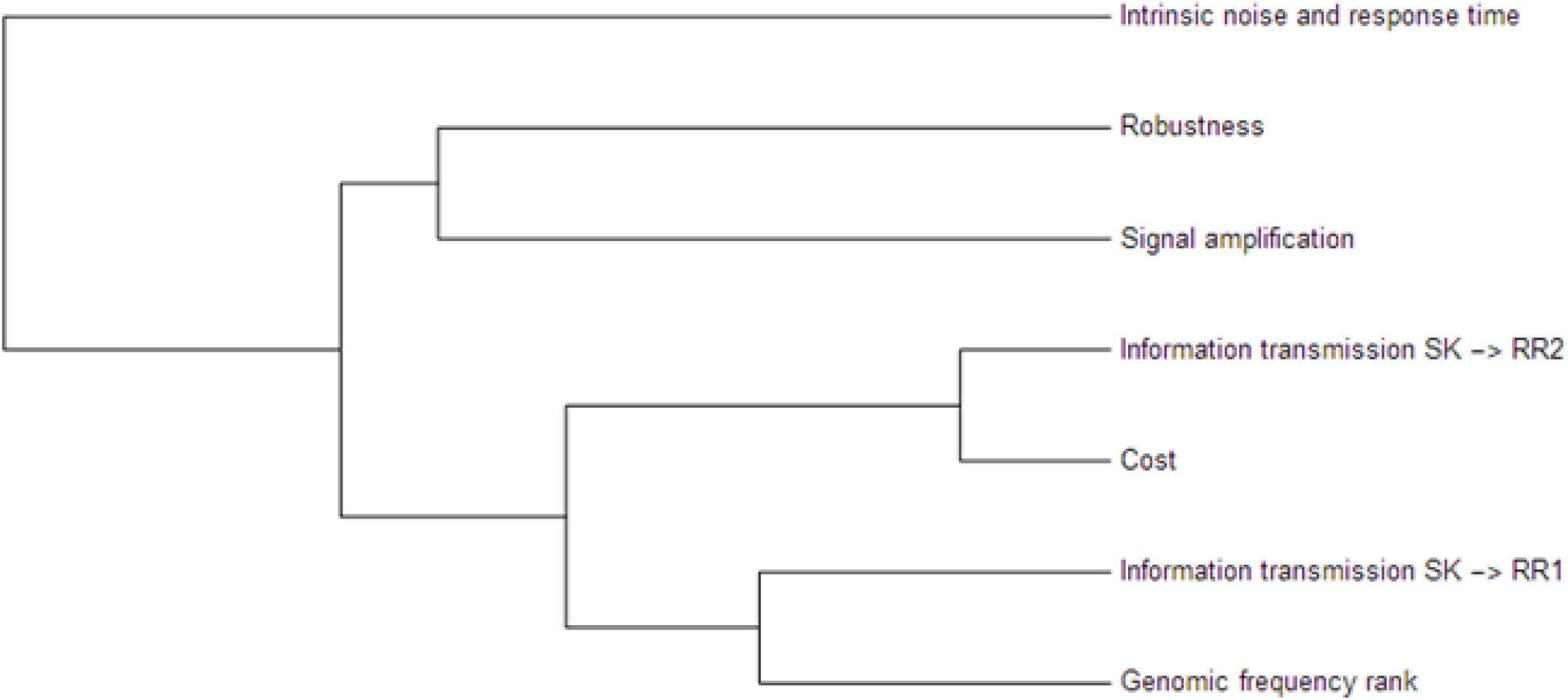
Clustering each architecture according to their abundance ranks in the ten percent best systems with respect to each of the physiological variables in Table 2. Variables that are absent in the clustering have an equal percentage of all architectures and therefore have equivalent ranking.

### Controlling for the possibility of bias in parameter space: data-driven comparative analysis of the five architectures

The previous results suggest that either cost of operating the circuit or information transmission or both could justify that architecture M3 is the most frequently found in genomic PRs. Still, given the percentage differences in architectural abundance between high performance and genomic PRs, it could be that we are analyzing parameter regions and protein abundances that are unexplored by nature, biasing our results.

To control for this possibility, we searched the literature for experimentally determined parameter values for the individual reactions of PR. A similar search was done for the total amount of the individual proteins in the circuit. Then, using a Latin hypercube approach (see methods) we created parameter values sets combining those values. Overall, five hundred combinations of parameters and concentrations were analyzed, percentages of abundance for each architecture in the set of systems that have the top 10% best performance were calculated as described above and clusters were built. The results are similar to those described before (Figure 2).

In order to clarify if cost or information transmission through the circuit are important factors in selecting PR architecture we performed two additional data driven sets of experiments.

### What physiological variables are optimized by native architectures in kinetically well characterized PRs?

First, we focus on two systems for which abundant quantitative and kinetic information is available: The Spo0 PR in *Bacillus subtilis* and the Sln1 system in *Saccharomyces cerevisiae*.

#### The Spo0 phosphorelay of *Bacillus subtilis*

We first focused on the Spo0 phosphorelay in *B. subtilis* because this is the PR system with the most abundant and reliable experimental determinations of kinetic information ^25–29^. The architecture of this system is of type M1, and supplementary Appendix S1 gives the protein abundances and parameter values for the reactions. We then created four alternative PR architectures (M2 - M4) with the same parameter values and protein abundances adjusted to minimize differences in dynamic behavior with respect to the native circuit architecture. Then, we compared the architectures with respect to the variables in Table 1. We found that the cost of protein synthesis is an order of magnitude lower for architecture M3 than for M1 (Figure 3). Protein costs for architectures M1, M2’, and M4 are an order of magnitude higher than in M3 and one order of magnitude lower than in M2. The robustness of signal transmission to environmental fluctuations in parameters is smaller in architectures M1 and M2’ and larger in M2, M3, and M4 (Figure 3). Architecture M4 is the fastest in responding to phosphorylating signals, and M4 and M1 are faster in a similar percentage of cases when the signal dephosphorylates the system (Figure 3). We clearly see that M1 transmits the highest amounts of information about the environment over the regulatable signal range for the Spo0 phosphorelay, whether the signal comes at the level of the SK (modulation of k1) or at the level of regulating dephosphorylation of the final response regulator (modulation of k26) (Figure 4). We note that the amount of information lost through the channel is smaller for architecture M2.

**Figure 3.**
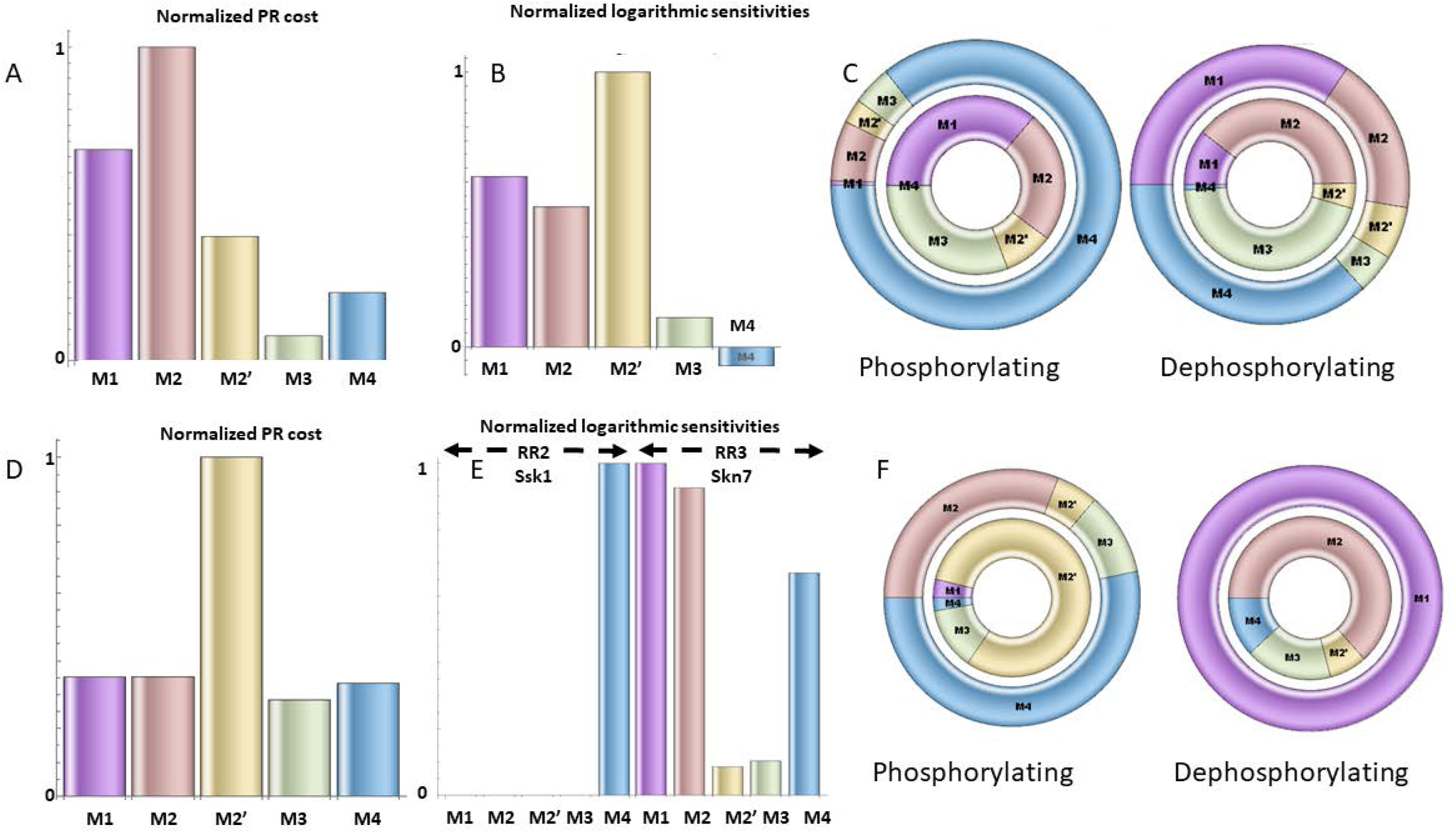
Effects of alternative architectures in the deterministic regime for the Spo0 phosphorelay of *Bacillus subtilis* and the Sln1 phosphorelay of *Saccharomyces cerevisiae*. In all cases, the protein amounts of the alternative architectures were optimized to make the steady state signal-response curves be as similar as possible (see methods). The black arrow indicates the native architecture. Panels A, B, and C pertain to the Spo0 phosphorelay. Panels D, E, and F pertain to the Sln1 phosphorelay. A, D – Cost of synthesizing the circuit under different architectures. X – axis: PR architecture. Y – axis: total metabolic cost of the circuit proteins (arbitrary units). B, E – normalized sensitivity of the steady state concentration of the final response regulator to changes in parameters. X – axis: PR architecture. Y -– axis: euclidean norm of the sensitivities vector of the response regulator. C, F – Percentage of transient responses where each architecture is the fastest (outer donut) or slowest (inner donut) in reaching 90% of the new steady state under equivalent conditions. Left-side donuts from responses to phosphorylating signals. Right-side donuts from responses to dephosphorylating signals.

**Figure 4.**
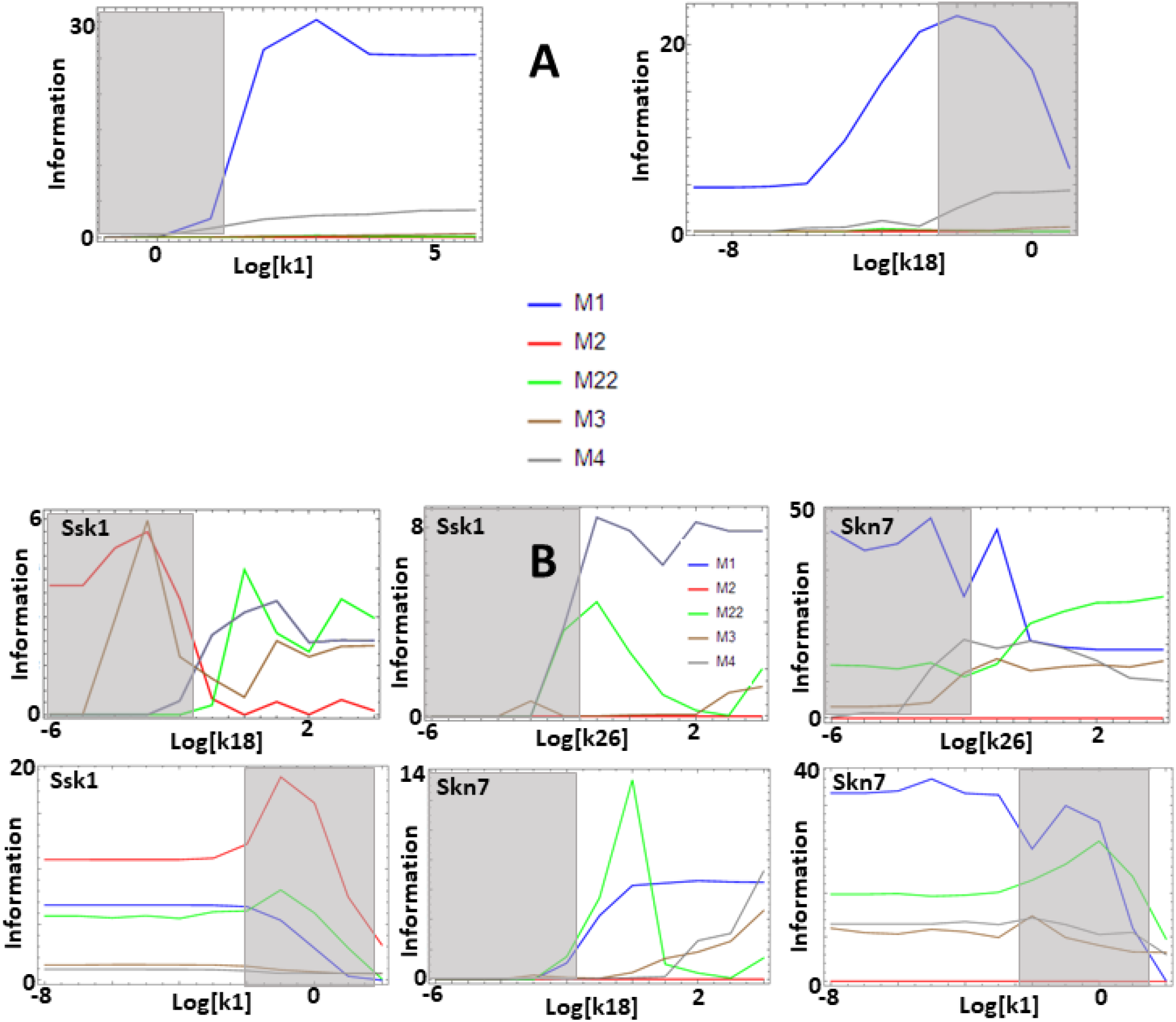
Effects of alternative architectures in the transmission of information through the Spo0 phosphorelay of *Bacillus subtilis* and the Sln1 phosphorelay of *Saccharomyces cerevisiae*. In all cases, the protein amounts of the alternative architectures were optimized to make the steady state signal-response curves be as similar as possible. **A** – Spo0 phosphorelay. Y-axis represents the accumulated mutual information over a range of 6 orders of magnitude for the self-dephosphorylation rate constant of kinA between variations in the number of phosphorylated kinA molecules and that of phosphorylated Spo0E molecules. k1 represents modulation of the kinA phosphorylation rate, while k18 represents modulation of SpoE dephosphorylation. Architecture M1 transmits the most information for comparable parameter values. **B** – Sln1 phosphorelay. Y-axis represents the accumulated mutual information over a range of 6 orders of magnitude for the self-dephosphorylation rate constant of Sln1 between variations in the number of phosphorylated Sln1 molecules and that of phosphorylated Ssk1 or Skn7 molecules. k1 represents modulation of the Sln1 phosphorylation rate, k18 represents modulation of Ssk1 dephosphorylation, and k26 represents modulation of Skn7 dephosphorylation. Architecture M2 transmits the most information to Ssk1 for comparable parameter values.

#### The Sln1/Ypd1/Ssk1 and Sln1/Ypd1/Skn7 phosphorelays of *Saccharomyces cerevisiae*

*S. cerevisiae* uses a PR with architecture M2 to sense changes in the osmolarity of the medium and regulate its internal metabolism and membrane composition, adapting to those changes. The hybrid sensor kinase Sln1 senses changes in membrane curvature and ultimately modulates the phosphorylation state of the terminal response regulators Ssk1 and Skn7. Phosphorylation of Ssk1 acts as a molecular switch in controlling the yeast’s osmosensing mitogen-activated protein (MAP) kinase cascade^30^, while phosphorylation of Skn7 directly affects the transcription of SLN1-SKN7-responsive genes ^31^. Skn7 is also involved in regulating heat shock response in a Sln1-Ypd1 independent way ^31^. Reliable kinetic parameter values and protein abundances are available for this PR.

We use those values to create a PR model where we consider the simultaneous presence of the terminal response regulators Ssk1 and Skn7. Then, create four independent alternative models, with architectures M1, M2’, M3, and M4 and parameter values equal to those of the native circuit architecture. Finally, we optimize protein abundance to minimize differences in the dynamic signal-response curve of each architecture with respect to that of the native architecture of the system.

We find that the cost of architecture M2’ is one order of magnitude higher than that of each of the other architectures, which have a similar metabolic cost (Figure 3). The sensitivity of the steady state concentrations for the Ssk1 RR are low and similar in all alternative architectures except M4, where sensitivity is high. In contrast, the sensitivity of Skn7 concentration is highest in architectures M1 and M2 (Figure 3). Architecture M4 most frequently responds faster to signals that increase the phosphorylation levels of the proteins, followed by M2 (Figure 3). In contrast, architecture M1 is always the fastest if the change in environmental conditions decreases phosphorylation levels of the circuit. When it comes to transmitting information about the environment to Skk1 over the regulatable range of the circuit, architecture M2 is the best for a wider range of parameters. It surprised us that all non-native architectures are better information transmitters to Skn7 than M2 (Figure 4). It is interesting to note that only very high rates of dephosphorylation for one RR affect the information transmitted by the circuit to the other, increasing it significantly.

Overall, these two systems suggest that transmission of information is an important determinant of architectural selection in PRs.

### Analyzing real PR examples: How do different PR architectures influence information transmission?

#### A performance atlas for information transmission in the parameter space of PR architectures

With the hypothesis that information transmission is important in the selection of PR architecture, we created a data-driven atlas that describes how architecture, protein abundance, and parameter values influence information transmission in PRs. To create that atlas we compiled experimentally determined parameter values and proteins abundances for as many PR systems as we could find in the primary literature and in public databases (see methods). Then we used those parameters and protein abundances to calculate the capacity to transmit information of each alternative architecture with all possible combinations of protein abundances and kinetic values. Finally we ranked the five architectures with respect to increasing capacity of information transmission. The atlas is summarized in Supplementary Table S2.

#### Experimentally determined protein abundances are consistent with amount of transmitted information being an important driver of PR architecture selection

We identified and obtained protein abundances for seventeen PR systems that are present in the PAXDB database and belong to seven different organisms^24^ (Table S3). Then we matched these abundances with the performance in the atlas of amount of transmitted information as a function of parameter values, protein abundance and circuit architecture given in Table S2. In 16 out of the 17 cases we find that the native architecture of the PR systems is consistent with maximizing the through-circuit transmitted information under experimentally known parameter conditions (Table 3).

**Table 3.**
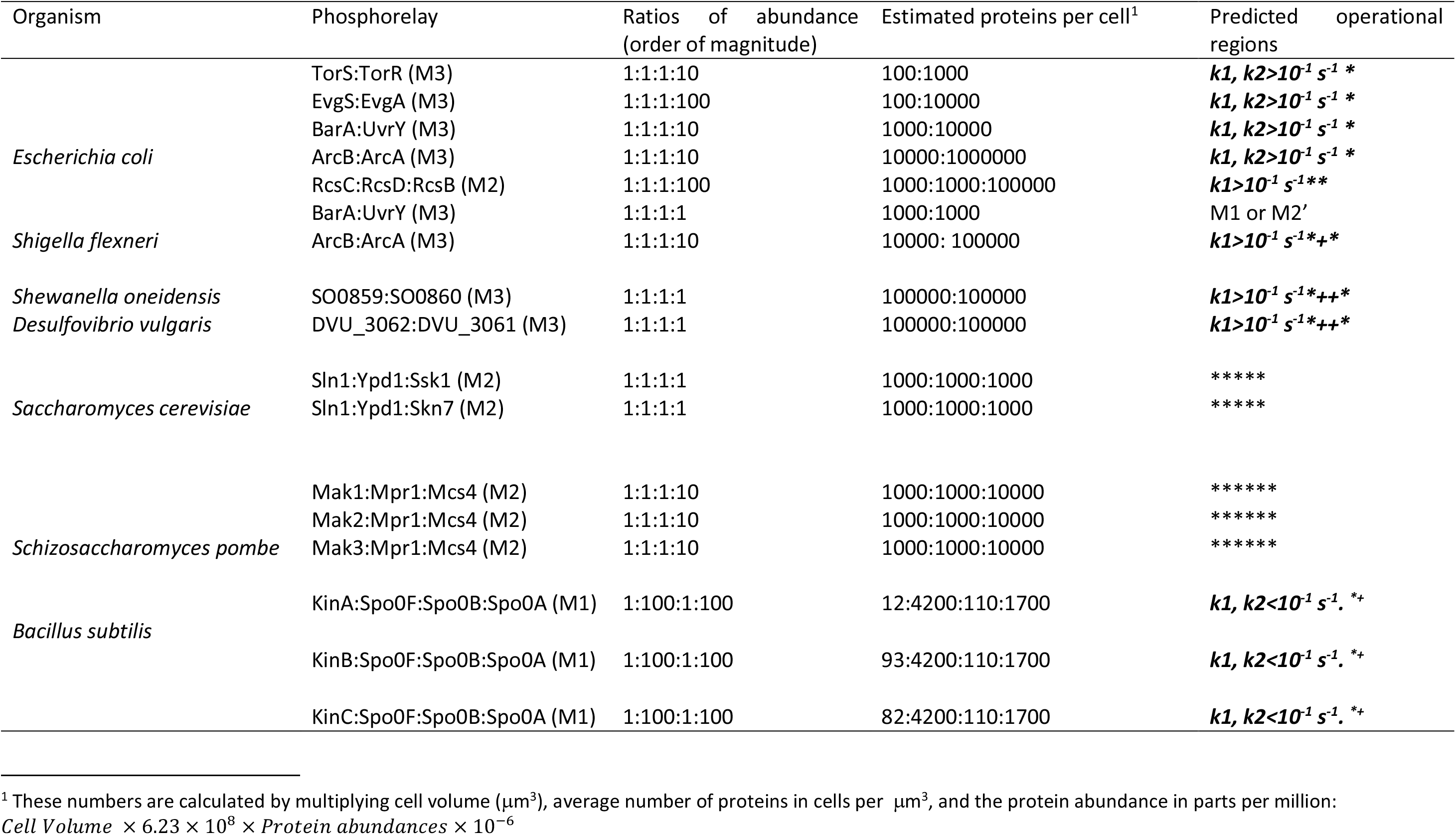
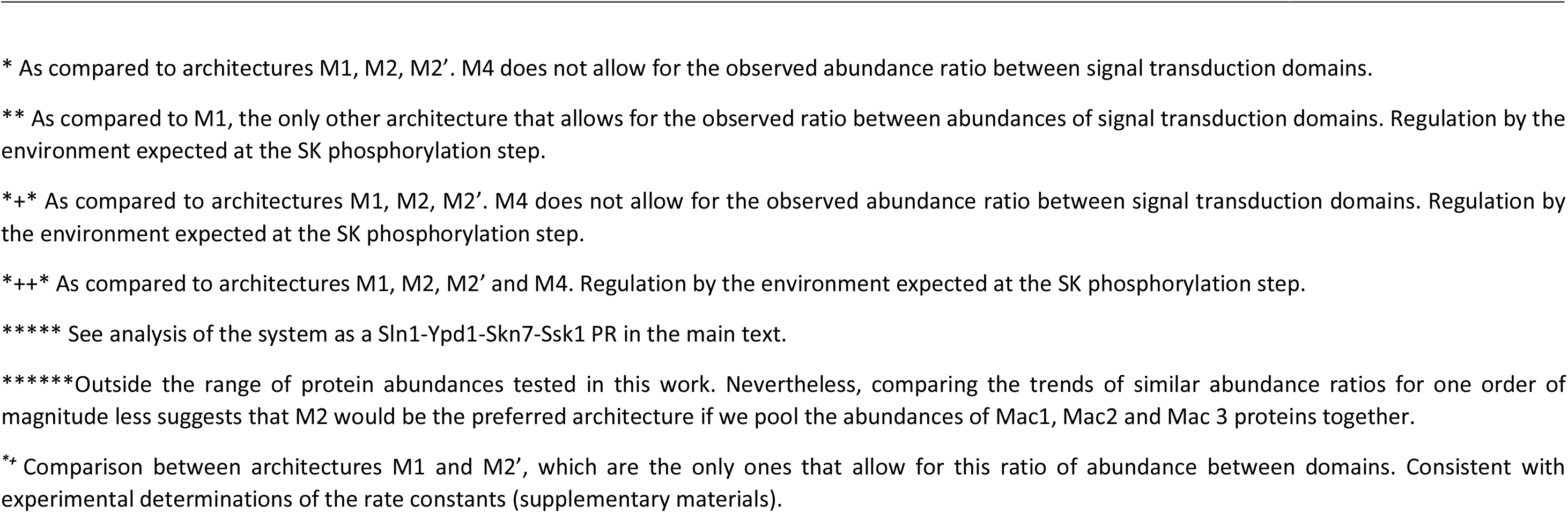
Observed native architectures and predictions of where in parameter space their observation is expected.

## Discussion

### Implications

Mutations in the genomes of organisms led to the emergence of variant architectures for biological circuits with similar biological function. Natural selection acting upon this variability led to fixing those architectures that are more efficient for the function of the circuit in the genome, contingent on life history. Elucidating the reasons why these architectures improve performance of the circuit reveals biological design principles that constrain evolution and explain the observed patterns.

In some cases, biological design principles are general and apply to a whole class of biological circuits. For example, overall feedback inhibition of the final product to the initial reaction of a linear biosynthetic pathway leads to molecular systems that are faster, more responsive, and less sensitive to fluctuations than any other regulatory alternatives ^32–34^. As such that regulatory solution is widespread in the tree of life. However, in many cases, design principles appear casuistic and are specific to a single circuit. For example, the circuit that regulates competence in *B. subtilis* appears to have been selected over alternative designs because of its noisy response ^35^, which enables bet edging in the competence response of the bacterium and improves its survival chances.

In the case of PR, a specific architecture is observed in over 72% of all genomic PR ^16^ in a sample of more than 9000 organisms ^4^. This suggested that indeed there could be design principles for selecting the architecture of these circuits. To test this hypothesis we needed to understand what physiological variables influence the performance of the circuit. Then, we needed to study how architecture affects those variables, allowing natural selection between alternative PR architectures.

Initially we compiled a list of physiological variables that are important for the function of these signal transduction circuits ^9,10,42–47,12,27,36–41^. We compared the alternative architectures to understand how they differed with respect to each of those variables. We then ranked the alternative designs with respect to their performance for each variable. Our results show that no unique variable could be used to rank all the alternative PR architectures in the same order as their genome abundance. Still, an analysis of large scale “simulomics” results shows that maximizing information transmission through the circuit and minimizing metabolic cost leads to an abundance of PR architectures that is the closest to the observed genomic frequency of those architectures.

This result led us to hypothesize that optimization of information transmission is an important driver of selection for PR architectures. To investigate this hypothesis we took a three step approach. First, we focused on two PR that are experimentally well characterized in terms of kinetic parameters and protein abundances. Simulation experiments revealed that the native architecture was the one that maximizes information transmission by the circuit. Second, we created a data-driven atlas of how architecture, protein abundance, and parameter values influence information transmission capacity. To create that atlas we compiled experimentally determined parameter values and proteins abundances for as many PR systems as we could find in the primary literature and in public databases. Then we used those parameters and protein abundances to calculate the capacity to transmit information of each alternative architecture with all possible combinations of protein abundances and kinetic values. Third, we took all PR for which experimental protein abundances are known and verified that the native architecture of the system is the one that maximizes information transmission in sixteen out of seventeen cases.

Overall, the results of our analysis for well characterized PR circuits are consistent with the hypothesis that information transmission is a crucial driver of PR circuit selection in nature. This was previously suggested to be the case for MAP kinase signaling and other metabolic circuits in eukaryotes ^48–55^. Still, architecture selection is a complex function of the interplay between the architecture, protein amounts and kinetic parameters, and evolution can play with these “genotypic variables” to find instances of each architectures that are almost equivalent information transmission circuits. Although we did look for additional specific examples of PR, we could not find any for which a complete set of parameter values had been measured either by the same or by independent labs. For some of them there were partial sets of parameter values (The ArcA – ArcB, the EvgS – EvgA, and the BvgS – BVgA PR). Nevertheless, creating models for these systems would have required using more than half of the parameters from other systems. This would have created a situation where the model would not be driven from the data, but rather from other estimates.

Our results provide evidence about design principles that could explain the evolutionary emergence of specific PR architectures. Those results also have consequences for synthetic biology. If we want to build synthetic PR that are stable in the genome, and have a significant level of quantitative understanding about the operating ranges for the circuit, then it should be designed using the architecture that has the highest capacity to transmit information over that range.

### Speculations

Our analysis invites some speculation regarding the ranges of operability for many PR. If 72% of all PR have a M3 architecture, as a first hypothesis we could expect that these systems operate in the ranges where the M3 architecture outperforms the others. By looking at our performance in information transmission atlas, this implies that M3 architecture systems are likely to be operating at abundance ranges of the PR that are below 100 molecules per cell for the protein containing the SK domain. This is amenable to testing in future proteomic experiments. The M3 circuits are also expected to be operating over modulatable ranges of the self-phosphorylation and self-dephosphorylation rate constants of SK above 10^−1^ min^−1^. Similarly, 18% of the identified PR might be operating mostly over the range where architecture M2’ transmits a high amount of information about the system, and 6% over the range where M4 is a better information transmitter.

In addition, we speculate that the type of environmental stimulus is also important in selecting the architecture. If the stimulus changes in a graded way and cells can be adjusted in a similar graded way, it makes sense that the architecture should allow for a high capacity transmission. For example, an architecture that allows the cell to distinguish between n+1 states (that is, with a capacity to transmit information of n+1 bits) provides for a better design that another architecture that only distinguishes between n states. On the other hand, if a very sharp response is required, architectures that can distinguish between a small number of states over a short operability range might be more effective.

Another tempting speculation arises from our modelling of the Sln1 phosphorelay. Given its information transmission profile, it could be that Sln1-mediated activation of Skn7 and its dependent genes may not be an important function of the circuit. For the experimentally determined parameter values and protein concentrations, other architectures would transmit more information to that RR. This suggests to us that Skn7’s role in heat shock response might be much more important for the cell than its role in osmoregulation.

### Limitations

Our study has several limitations. Here we discuss those we think are the most important. First, we may have underestimated the percentage of PR with architecture M1, as these PR may be coded under different promoters in separate parts of the genome. This could also be true for designs M2, M2’, and M3. Nevertheless, the amount of proteins of type SKRR, SKHpt, or SKRRHpt that are found without the remaining operon in genomes is much smaller than that of the same type of proteins that are assembled in operons with the remaining putative cognate proteins of the PR ^4^, which indicates that any underestimation of these architectures is likely to be small.

Second, the performance landscape of PR can be strongly affected by how its expression is regulated ^45,56–58^. Nevertheless, transcriptional regulation occurs on a timescale of tens to hundreds of minutes, while the regulation of phosphorylation levels of the PR occurs on a timescale of minutes and this timescale can also strongly influence the performance of the gene expression regulatory circuit ^57,59^.

Third, there might be a wider range of parameter values for each reaction of the PR in the wild and this could change our phenotypical mapping of PR behavior onto the phase and architectural spaces of the circuits. Nevertheless, if this is so, our results would still be valid for the regions that were analyzed and considering that expanded set of values would only increase the “genotypic space” without changing the mapping we present here.

Fourth, our analysis of the capacity to transmit information focuses on steady state shifts. One could argue that the transient capacity could be a more important determinant of system performance. However, we tested how the transient capacity to transmit information compares to the corresponding steady state capacity and found that the latter is always a majorant of the former.

## Methods

### Mathematical Modelling

PR shown in Figure 1 represent more than 99% of all PR circuits present in the fully sequenced genomes of more than 9000 organisms. This information was obtained from ^4^. Two types of models were created for each architecture.

#### S-system and GMA models

We used the power law formalism for the large-scale mapping of dynamic behavior into parameter and concentration space. This formalism can be used to create quantitative models of circuits when little kinetic and/or mechanistic information is available ^60^.

First, we created models for the five architectures using the S-system representation. This representation allows us to calculate analytical expressions for the steady state values of each of the deterministic physiological variables that we needed to evaluate. Then we used the generalized mass action representation to create models that could be evaluated numerically for each of the five architectures. These models were used to map the physiological behavior of each architecture in parameter space. All models are given in supplementary Appendix S1.

#### Mechanistic Models

We used a mass action description of reactions to create a mechanistically more detailed model for each architecture ^60^. These models are given in supplementary Appendix S1. They were parameterized with experimentally determined parameter values and protein concentrations compiled from the primary literature. We used these models to create an atlas of performance with information transmission through the PR as a function of protein amount, parameter values, and circuit architecture.

#### Estimating protein concentrations

Proteins occupy approximately 20% of cell volume across the tree of life ^61,62^. Taking into account average protein sizes, average protein masses and factoring in cell volumes one can estimate that total protein concentration in cells is of the order of ~ 1 μM (ranging between 0.4-1.4) ^63^, or 6.23 × 10^6^ proteins per μm^3^. We then use the average cell volume for the different cell types to estimate the total amount of proteins per cell.

In order to estimate biologically relevant protein amounts for each circuit topology we used whole-proteome relative abundance determinations (Table S3) reported by PaxDB ^24^. The database information is given in parts per million. By multiplying these number by total number of cell proteins, we calculate how many proteins exist for each experimental PR system, within the experimental error.

With this information we can further estimate typical orders of magnitude for the ratios between protein abundances within a PR circuit. In general, we find that protein abundances and ratios between protein abundances are bound and have a limited range in PR (Table S3).

#### Parameter values for the mass action models

To find experimentally determined parameter values for the individual reactions of the PR, we searched Medline and the Biomodels database ^64^. We searched for previous mathematical models for TCS and signal transduction PR. In addition we also search the primary literature for known circuits that have been characterized biochemically and collected all different parameter values for the various reactions in the PR circuit. This information is compiled in Supplementary Table S4, revealing that kinetic parameters for corresponding reactions in experimentally well-characterized systems are quite similar and, in most cases, within the same order of magnitude.

### Calculating the dynamic behavior of each model

#### Steady state concentrations and stability

We obtained steady state concentrations by numerically solving the GMA and mass action models for each set of parameter values. We then calculated the jacobian of the ODE system and the eigenvalues of that jacobian to determine the stability of the steady state. We compared the steady state stability of the PR circuits in two ways. As we selected for sets of parameter values that generate systems with stable steady states (see methods), all real parts of the eigenvalues of the steady state are negative. Taking this into account we compared the minimum of the real parts of the eigenvalues in each pair of models. This comparison allows us to compare the fastest time scale in which the two circuits respond to a transient perturbation to their steady states. We also compared the maximum of the real parts of the eigenvalues between the two circuits in each pair. This allows us to compare the slowest time scale in which the two circuits respond to a transient perturbation to their steady states.

#### Logarithmic gains and parameter sensitivities

To estimate the response of a physiological variable to changes in environmental signals or model parameters, we compute ^60^

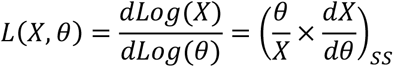

Here, ss indicates evaluation at a reference steady state.

#### Signal amplification

To estimate signal amplification we calculated the logarithmic gains of the concentration of the phosphorylated form of the final response regulator (RR_2_) with respect to the cognate signal of the system (Signal), as given by:

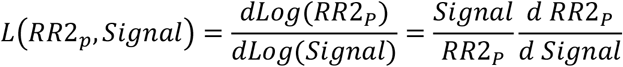

#### Sensitivity to the total amount of protein

To estimate sensitivity of the different phosphorylated forms of the PR proteins (Prot-P) with respect to the total amount of circuit proteins (Pr_tot_), we calculated the logarithmic sensitivities, as given by:

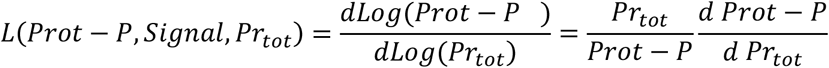

#### Global sensitivity to fluctuations in parameters

To estimate sensitivity of the different phosphorylated forms of the PR proteins (Prot-P) with respect to fluctuations in each of the parameters (Par_i_), we calculated the logarithmic sensitivities, as given by:

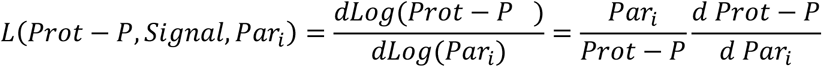

Then, for each protein, we calculate the norm of the vector whose coordinates are the individual sensitivities of that protein, as described in ^60^. This is an aggregated measure of the robustness of circuits to fluctuations in parameter values ^47,65^.

#### Metabolic cost of the circuit

To calculate the metabolic cost of running a circuit we focused on two aspects. First we estimated the average size of each type of protein in each architecture from the data in ^4^. Then, we multiplied the number of amino acids of the individual proteins in the phosphorelay by the number of proteins in the cell and used this number as a proxy for the cost of each circuit. Then, we also focused on the reactions of the circuit that consume ATP (self-phosphorylation of the SK domain).

#### Response times

We performed two independent experiments per architecture and per set of parameter values to calculate the response times. First, we start with the system fully dephosphorylated and run a time course simulation using the same scan procedure for the self-phosphorylation and self-dephosphorylation rate constant of SK. We then calculate the time it takes each of the architectures to reach 90% of the new steady state values. We repeated this experiment starting with fully phosphorylated proteins.

#### Steady state noise: the coefficient of variation

To estimate the intrinsic noise for each architecture and set of parameter values we numerically calculated the deterministic concentration of the systems, converted this concentration into number of copies of the PR, and ran twenty five stochastic simulations of twenty minutes. Then, for each variable we calculated the coefficient of variation (CV) as given by:

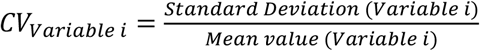

#### Amount of information transmitted through the circuit

To calculate the information transmitted through the PR we simulated that system to steady state using Gillespie’s algorithm for stochastic simulation one hundred times per set of parameter values ^66^. The proxy for changes in the environmental information was taken to be the number of phosphorylated molecules of the SK domain, as this is the sensing domain of the PR. The output of the PR was considered to be the number of phosphorylated final response regulators. The information transmission through the circuit was estimated using the standard formula for mutual information between the two variables:

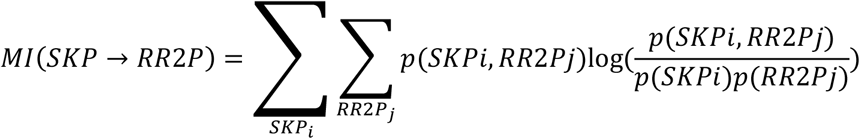

The information transmission from the SK to the first RR was estimated from:

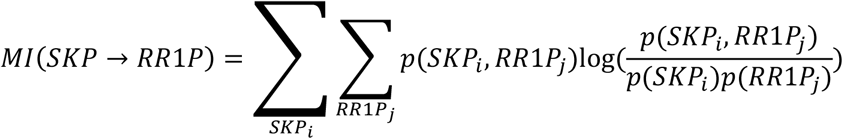

The information transmission from the Hpt domain to the final RR was estimated from:

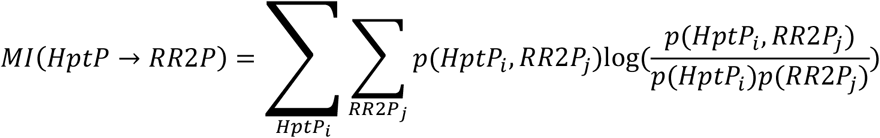

In these formulas, p(x) is the relative frequency of x, and p(x,y) is the joint relative frequency of x and y.

We compare the information transmitted through the various steps of the circuit as a measure of information loss. The closer *MI*(*SKP* → *RR*1*P*), *MI*(*RR*1*P* → *HptP*), and *MI*(*HptP* → *RR*2*P*) are, the smaller the amount of lost information.

### Comparison of dynamic properties between architectures

The comparison between alternative architectures was done in a pairwise manner, to facilitate interpretation.

#### Analytical comparison

We calculated the analytical solutions for the S-system model of each architecture. Then we took the ratio of corresponding properties between all possible pairs of architectures, controlling the comparison in such a way that the phosphorylated levels of the final RR were the same in both elements of the pair. If the ratios could be demonstrated to be always larger or smaller than one, this would mean that the property being analyzed would always be large in one of the architectures.

#### Numerical comparison

We took a reference architecture and generate a set of parameter values drawing from a random distribution of parameters with realistic boundaries. We did the same sampling for protein concentrations. Then we calculated the numerical properties of the pair of models being compared, discarding all sets of parameter values that led to systems with negative signal amplifications, unstable steady states, aggregated sensitivities to fluctuations in parameters and sensitivities of phosphorylated forms to total amounts of circuit protein that were too high, and response times that were too slow. We stopped generating sets of parameter values when 5000 of those set met all criteria of realist performance described above. Overall, we tested several hundred million sets of parameter values in all comparisons.

### Mathematical modelling of the Spo0 and Sln1 phosphorelays

Detailed mechanistic models were created for the Spo0 and Sln1 phosphorelays, as described in supplementary Appendix S1. These models use experimental data from ^29^ for Spo0 and from the SGD ^67^ for Sln1.

Then, for each of the PR, four additional equivalent models were created, each assuming that the PR would have a different architecture. The parameter values for these alternative architectures were considered to be the same as those for the native circuit architecture. The concentrations for the proteins were optimized by first simulating the original model at different signal intensities and calculating the steady state phosphorylation state of the final RR. Using these results we then performed the same experiments for each alternative architecture and allowed for the proteins that got fused or separated with respect to the cognate architecture to change in concentration by up to one order of magnitude below or above that in the natural architecture. We then selected the concentrations that led to the most similar signal-response curves using a least square minimum criteria for the differences between phosphorylated final RR.

### Calculating the cost of alternative architectures for the Spo0 and Sln1 phosphorelays

To calculate the cost of alternative architectures for the Spo0 and Sln1 we first counted the number of amino acids in the sequence for each protein in the original architecture. Then, we assumed that any alternative architecture would result from fusing or splitting the original proteins while conserving the same number of amino acids. Finally, we multiplied the number of amino acids of the individual proteins in the phosphorelay by the number of proteins in the cell and used this number as a proxy for the cost of each architecture.

### Software

All models and analysis were done using Mathematica ^68^. The notebooks containing all code to generate each figure are given as supplementary data pack S1.

---

**Table S1.**
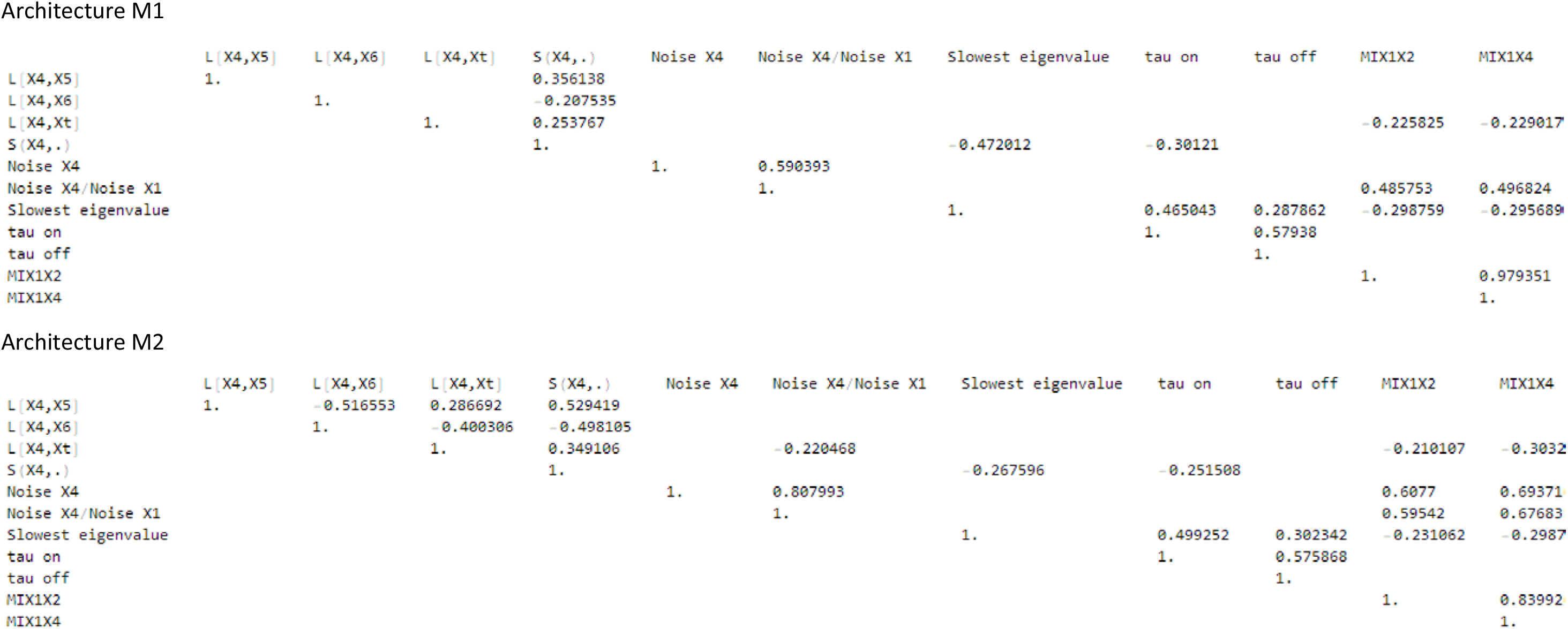
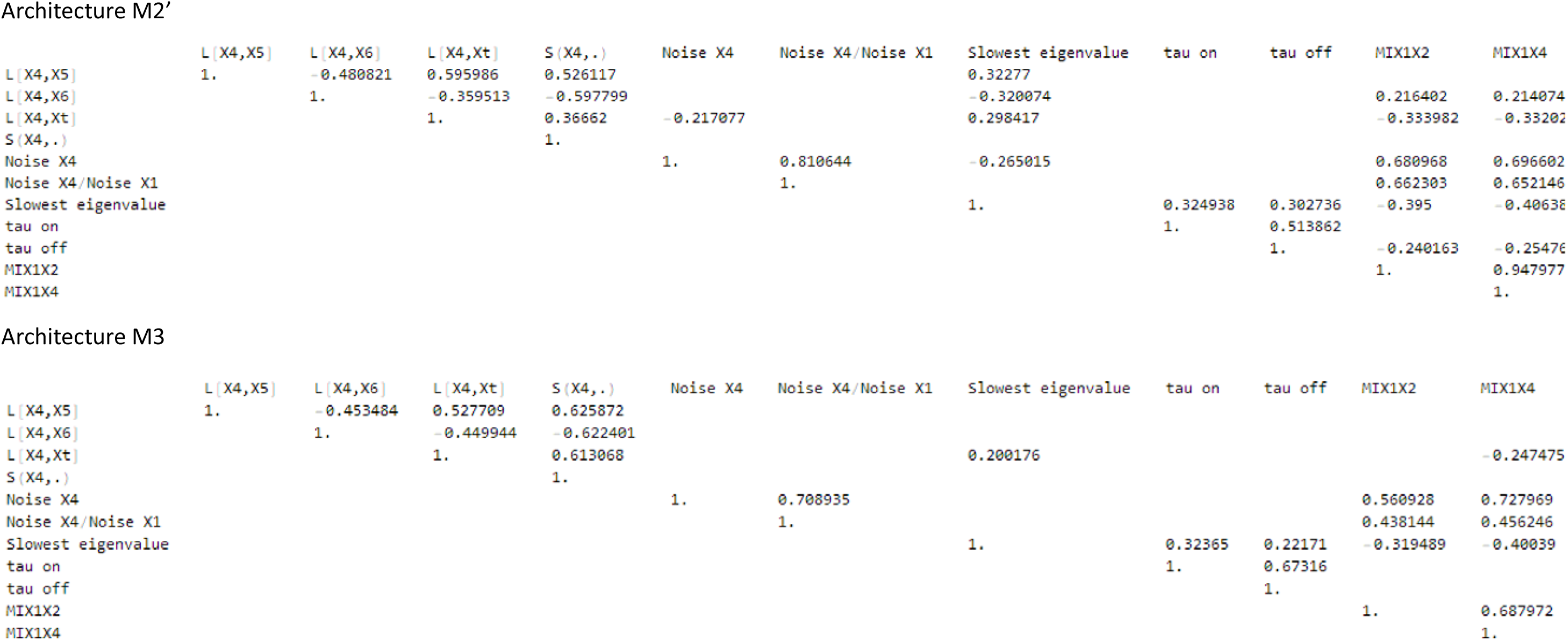
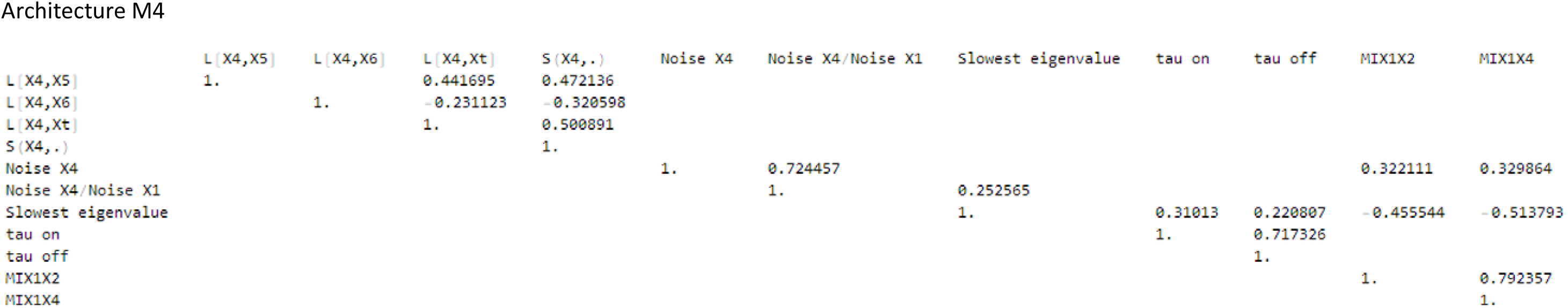
Correlation analysis of variable in different architectures: Spearman correlation.

**Table S2 – Performance atlas with respect to information transmission through the phosphorelay as a function of architecture, protein amounts, and parameter values**

**Table S3.**
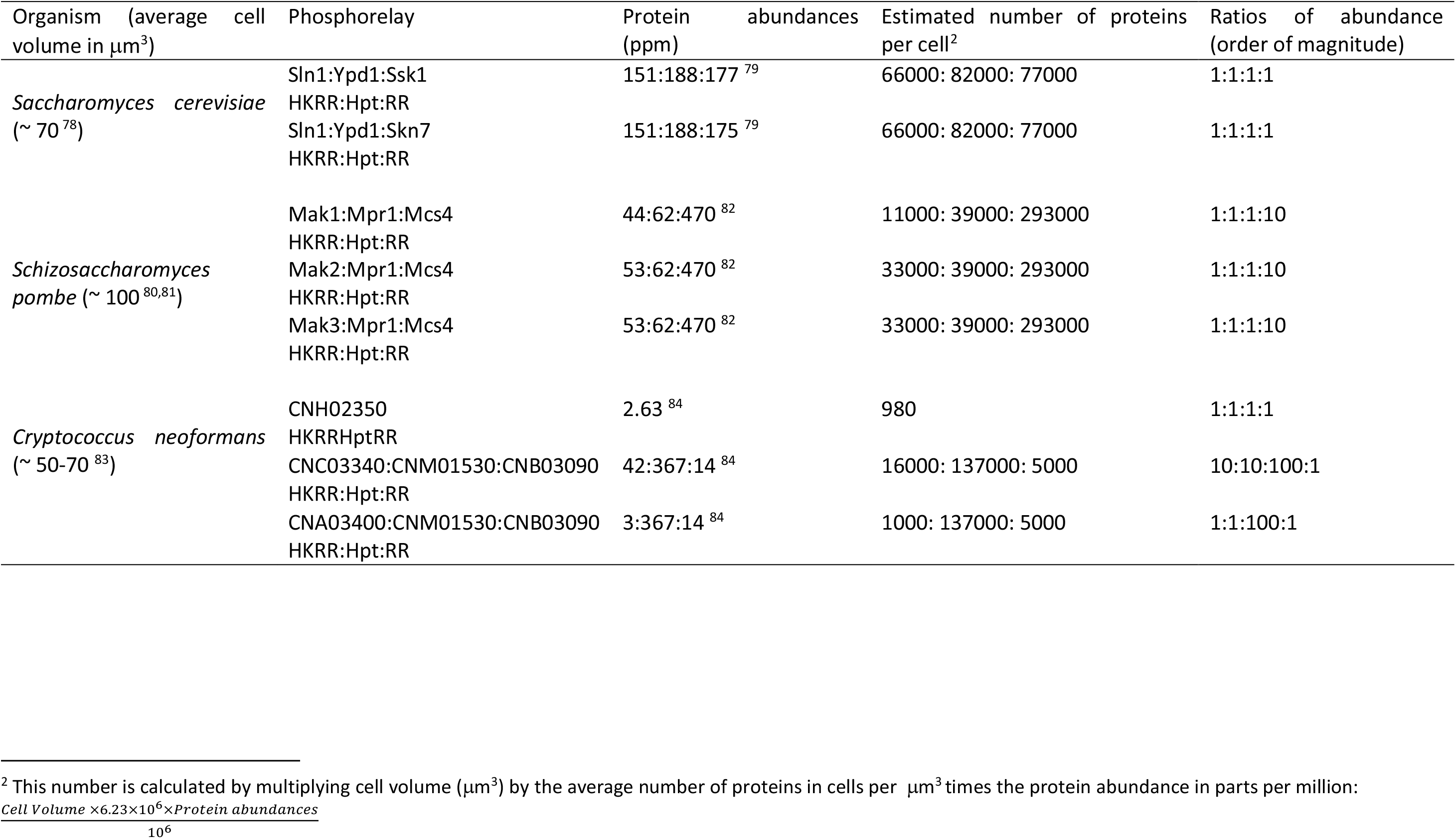
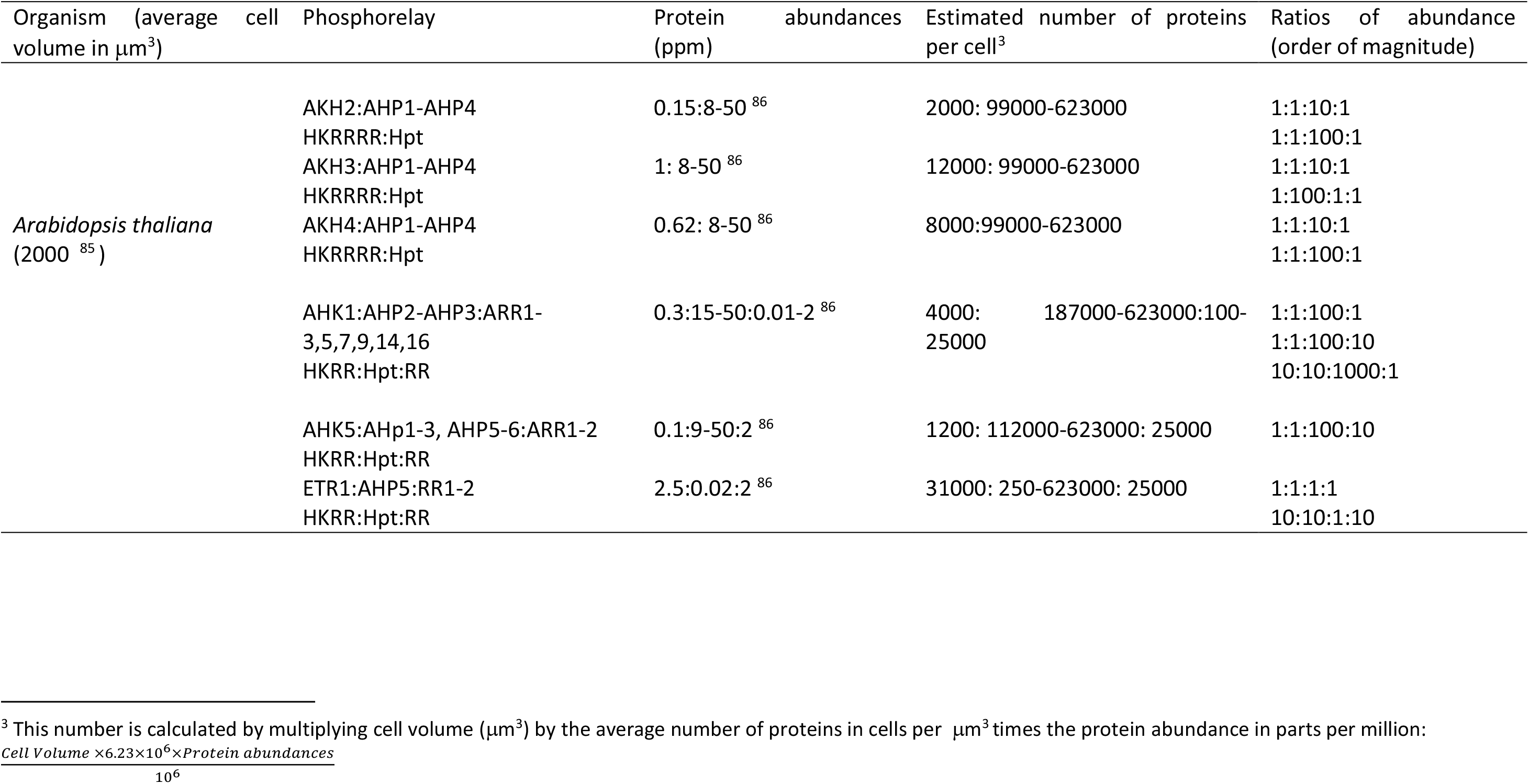
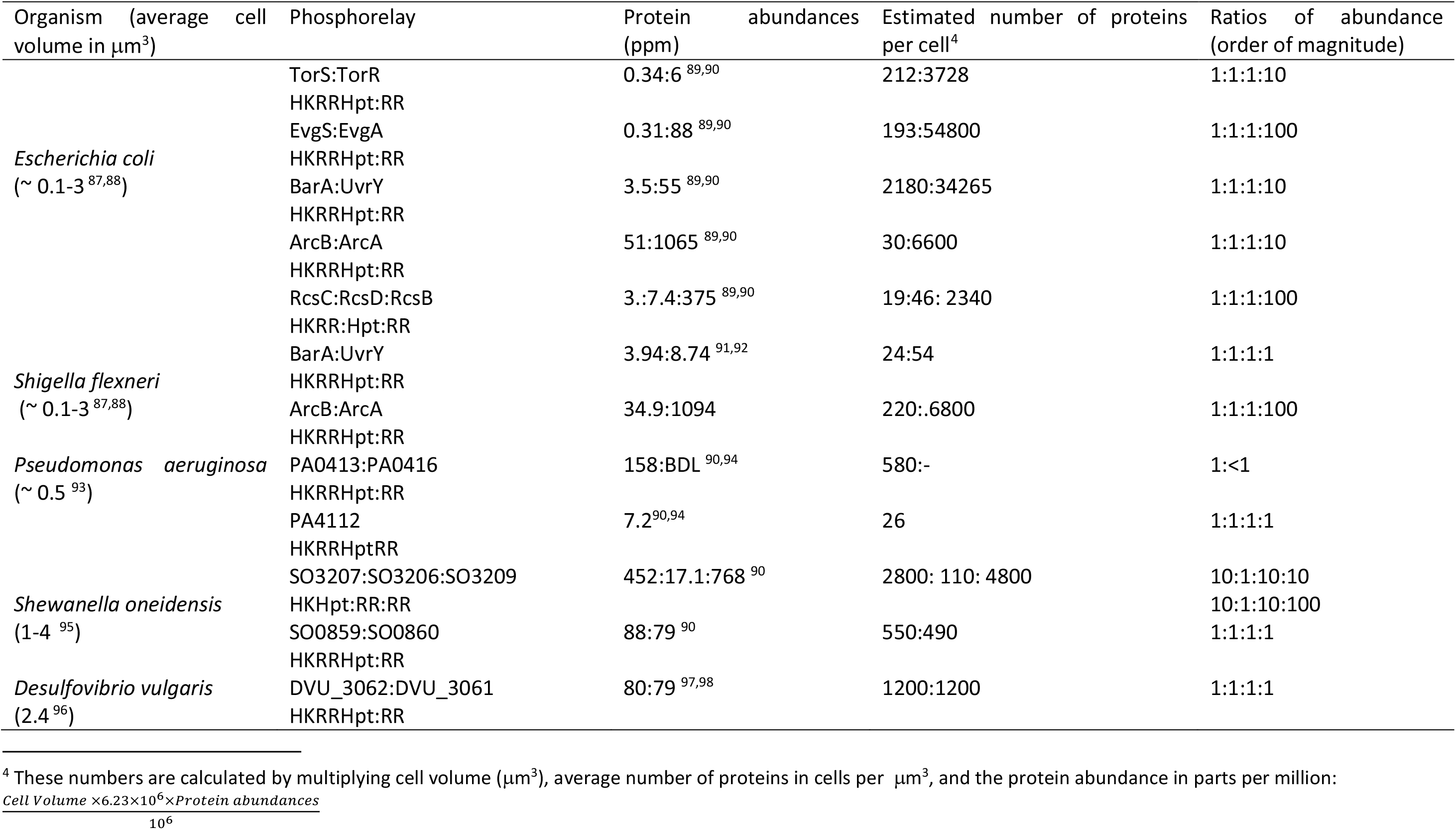
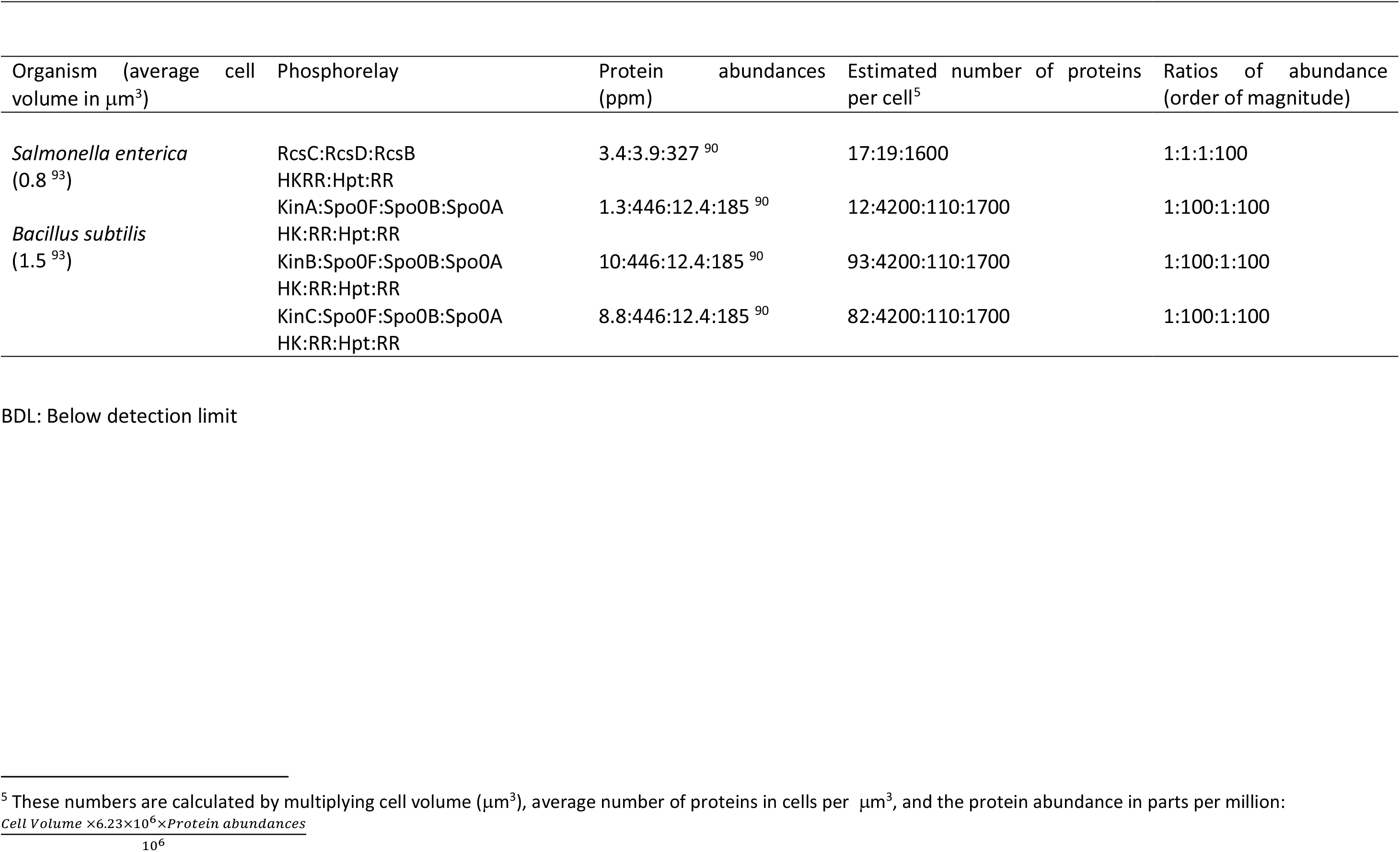
Phosphorelay circuits for which there is abundance information.

**Table S4.**
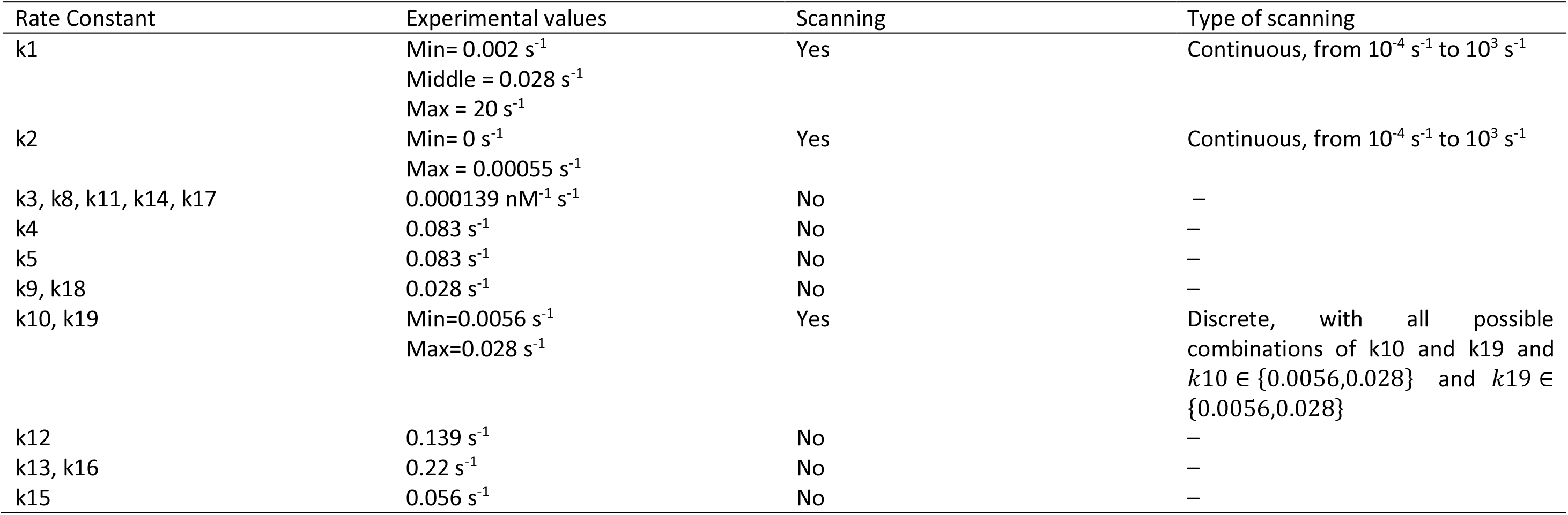
Numerical values of the rate constants of the 19 reactions in the PR models used in the stochastic simulations, represented in Figure 4. These values are based on ^29^. We found a range of variation in four of the rate constants (k1, k2, k10 and k19) ^25–27,43^. For the generality of the results, we used both the minimum and the maximum value of these four rate constants, obtaining 16 different combinations of numerical values. Each one of these 16 combinations of rate constant values was used in each one of the 5 PR models to obtain 100 stochastic trajectories.

| Rate Constant | Experimental values | Scanning | Type of scanning |
| --- | --- | --- | --- |
| k1 | Min= 0.002 s<sup>-1</sup><br>Middle = 0.028 s<sup>-1</sup><br>Max = 20 s<sup>-1</sup> | Yes | Continuous, from 10<sup>-4</sup> s<sup>-1</sup> to 10<sup>3</sup> s<sup>-1</sup> |
| k2 | Min= 0 s<sup>-1</sup><br>Max = 0.00055 s<sup>-1</sup> | Yes | Continuous, from 10<sup>-4</sup> s<sup>-1</sup> to 10<sup>3</sup> s<sup>-1</sup> |
| k3, k8, k11, k14, k17 | 0.000139 nM<sup>-1</sup> s<sup>-1</sup> | No | – |
| k4 | 0.083 s<sup>-1</sup> | No | – |
| k5 | 0.083 s<sup>-1</sup> | No | – |
| k9, k18 | 0.028 s<sup>-1</sup> | No | – |
| k10, k19 | Min=0.0056 s<sup>-1</sup><br>Max=0.028 s<sup>-1</sup> | Yes | Discrete, with all possible combinations of k10 and k19 and $k10 \in \{0.0056, 0.028\}$ and $k19 \in \{0.0056, 0.028\}$ |
| k12 | 0.139 s<sup>-1</sup> | No | – |
| k13, k16 | 0.22 s<sup>-1</sup> | No | – |
| k15 | 0.056 s<sup>-1</sup> | No | – |

## Supplementary text S1 - Mathematical modelling

### Power law models

Architecture M1

The conceptual scheme we used to derive the power law mathematical models for architecture M1 is as follows

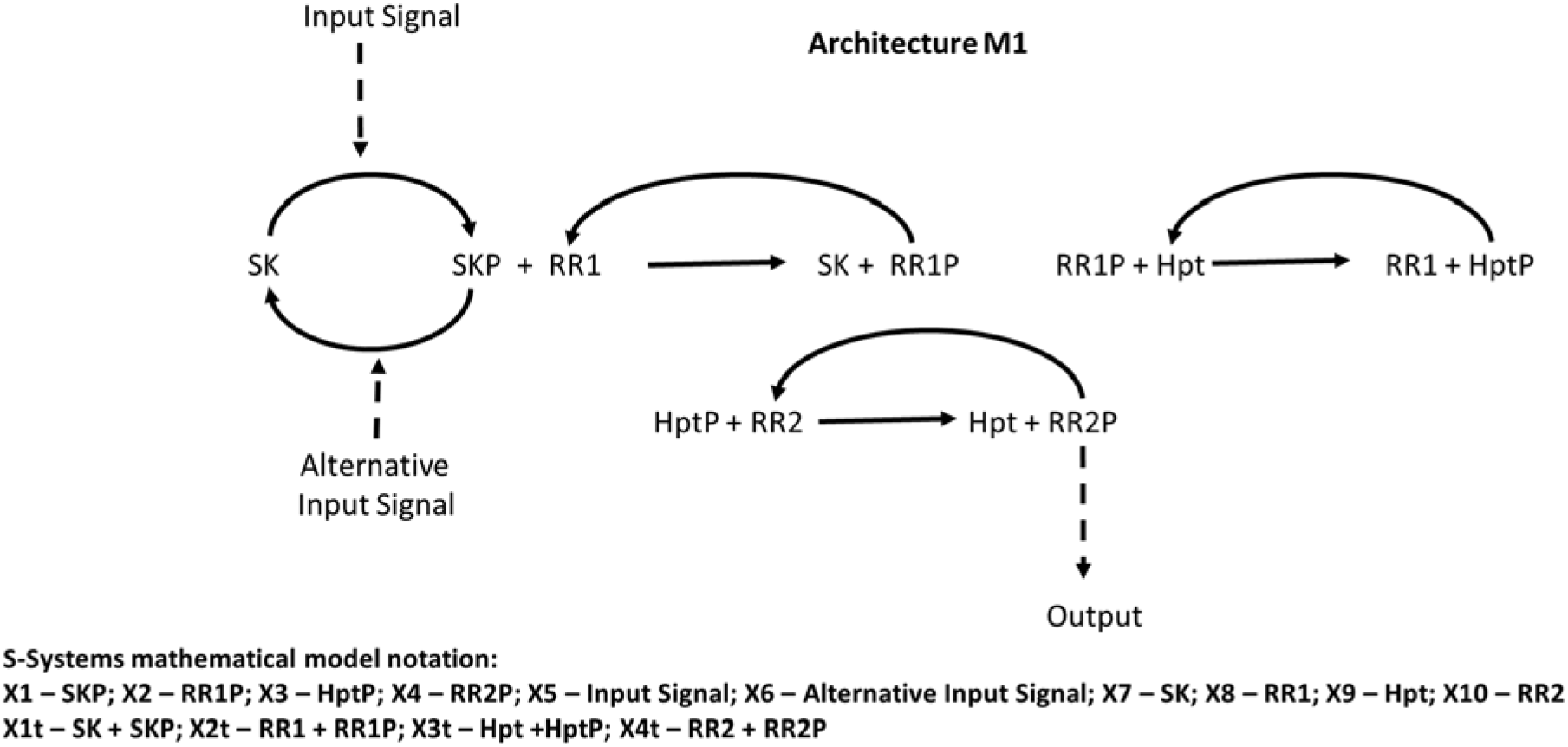

GMA Model

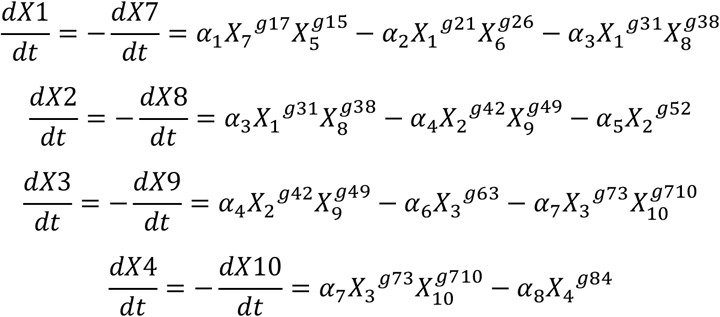

S-system Model

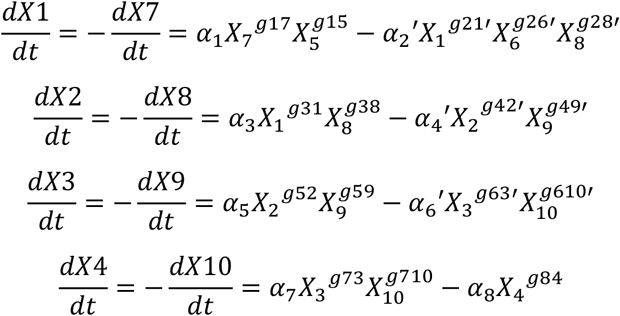

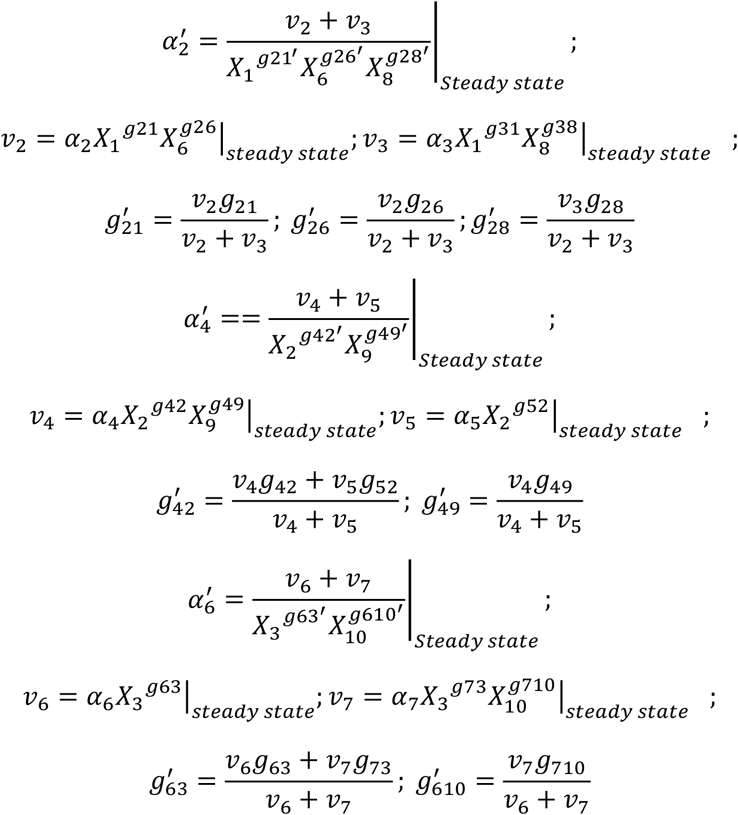

Architecture M2

The conceptual scheme we used to derive the power law mathematical models for architecture M2 is as follows

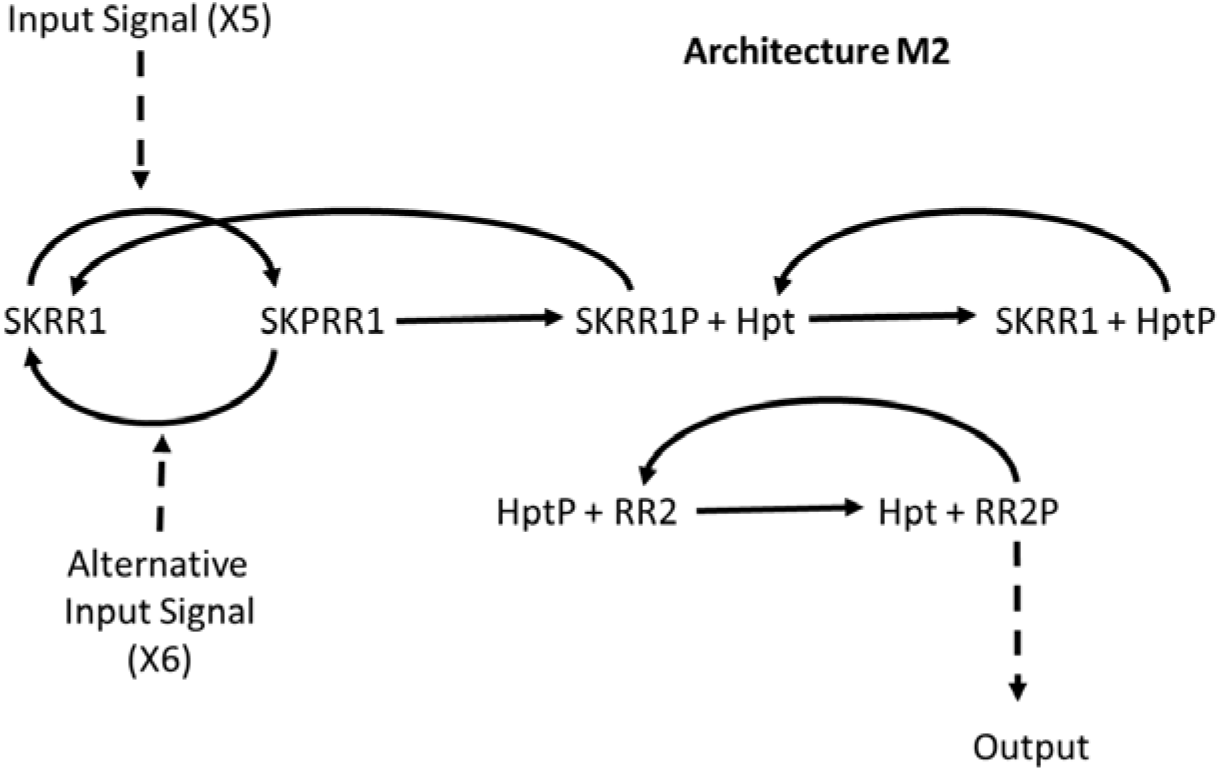

GMA Model

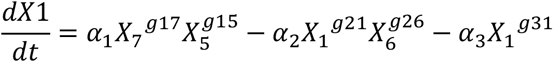

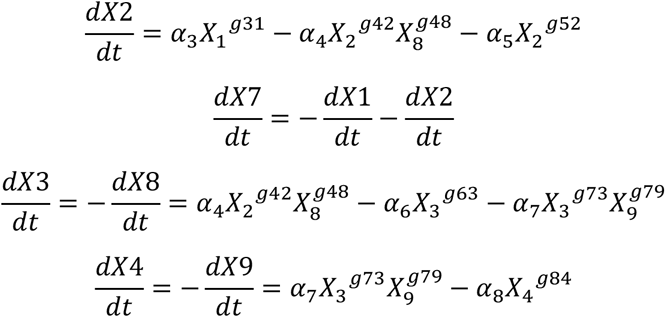

S-system Model

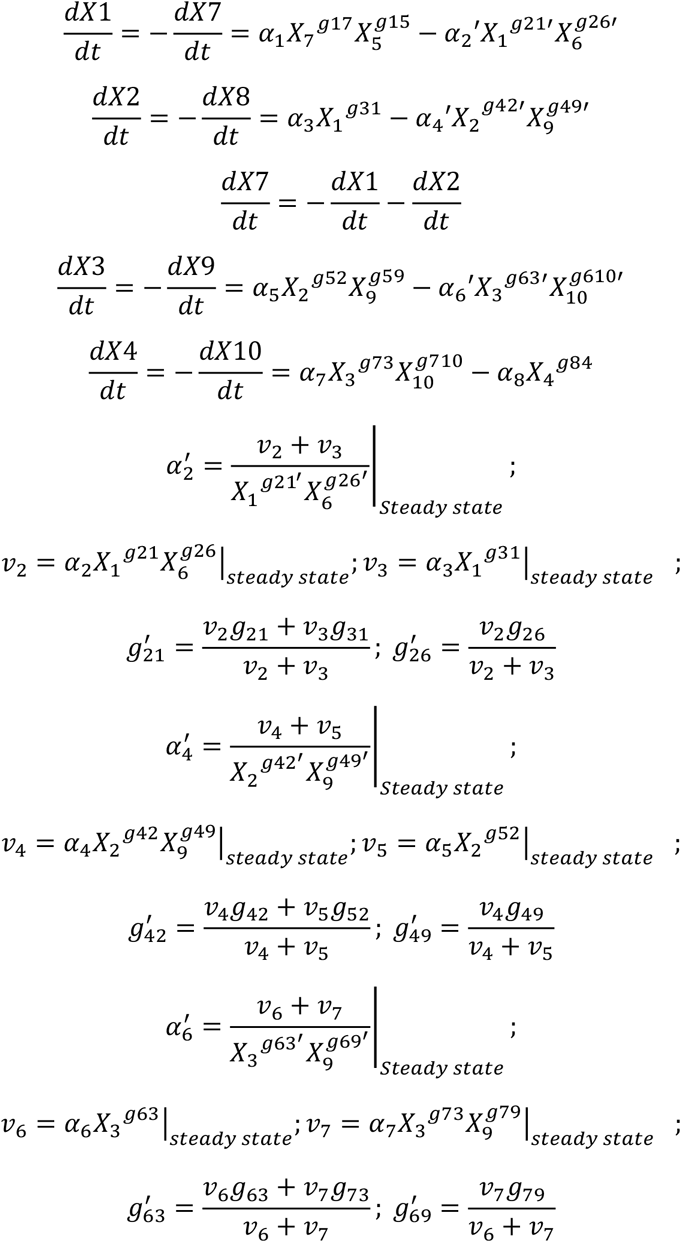

Architecture M2’

The conceptual scheme we used to derive the power law mathematical models for architecture M2’ is as follows

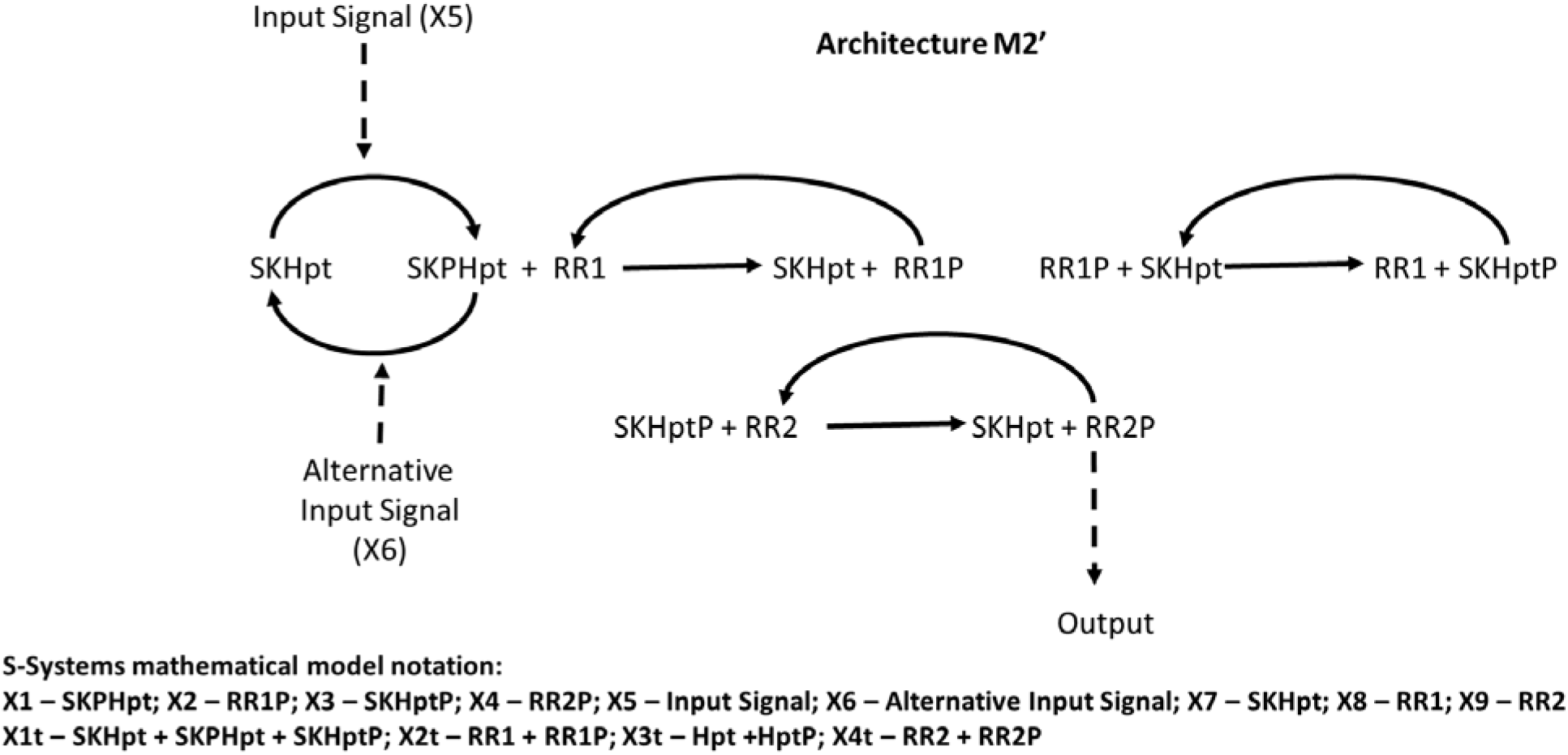

GMA Model

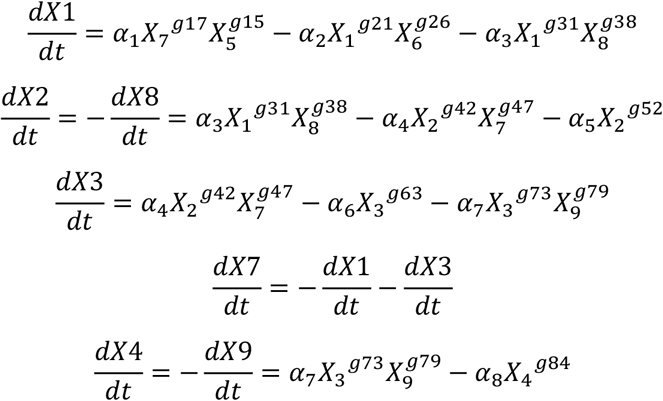

S-system Model

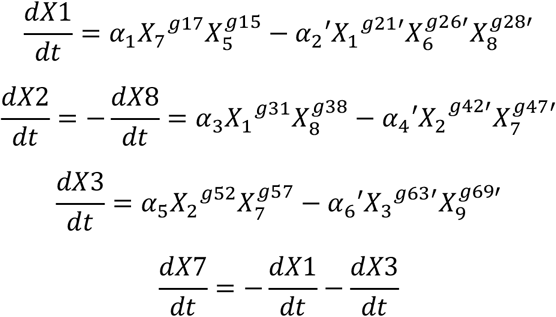

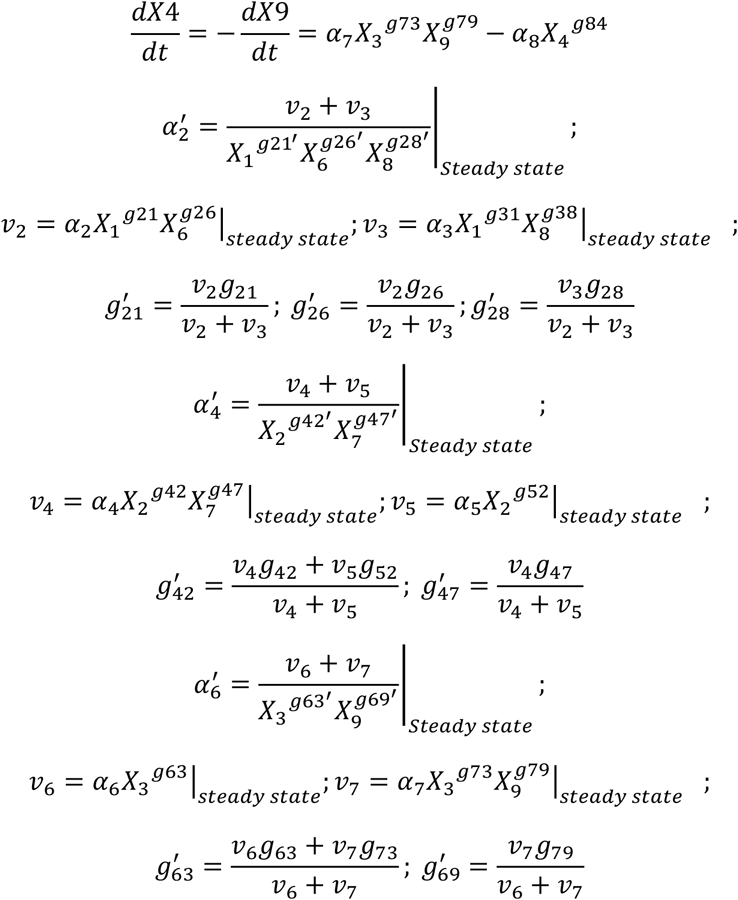

Architecture M3

The conceptual scheme we used to derive the power law mathematical models for architecture M3 is as follows

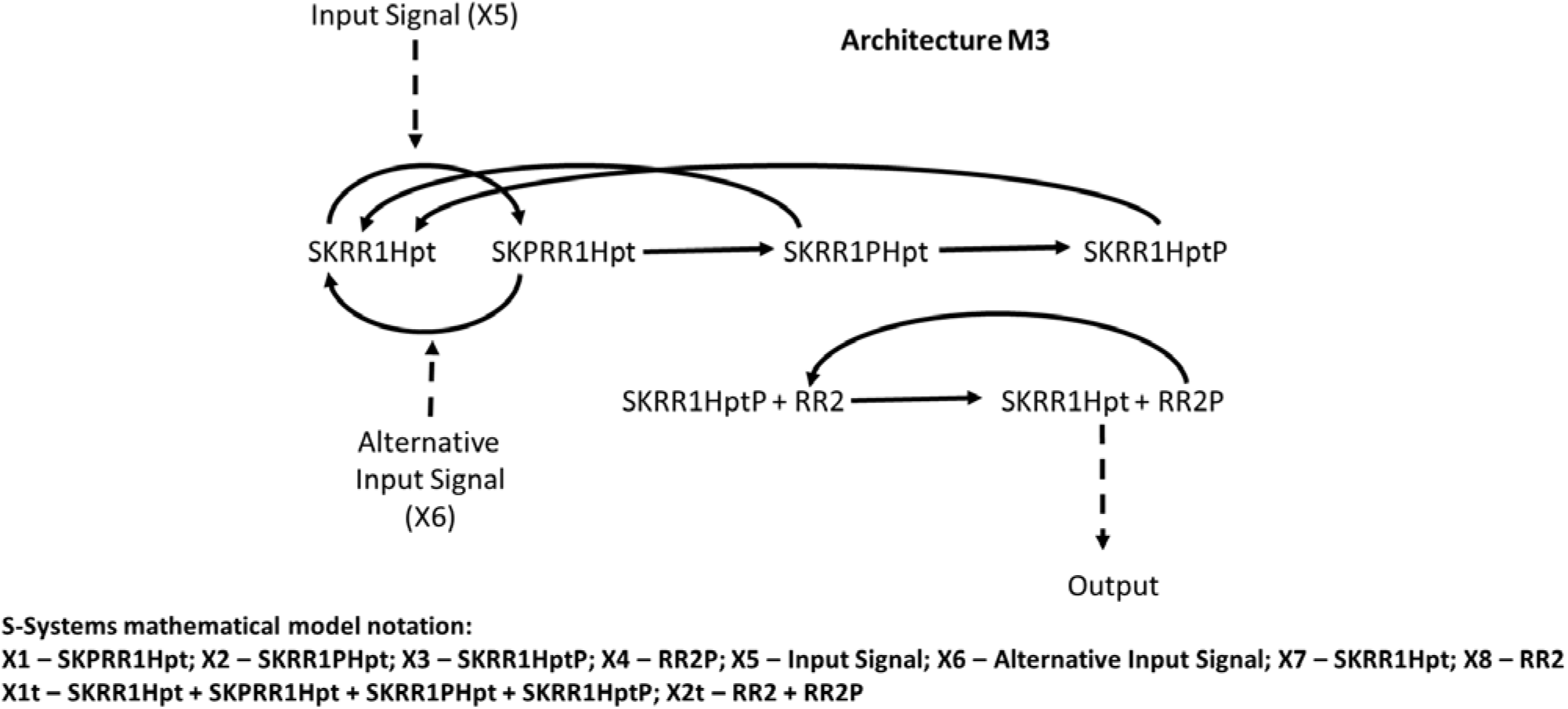

GMA Model

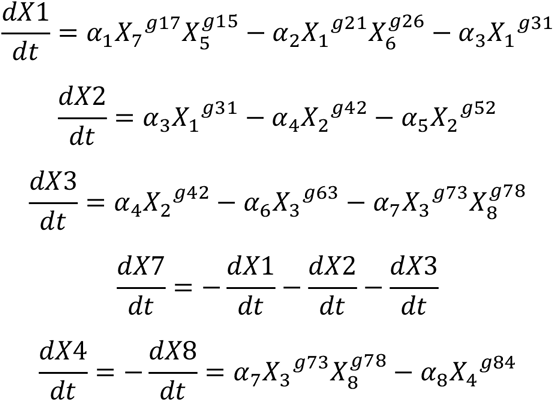

S-system Model

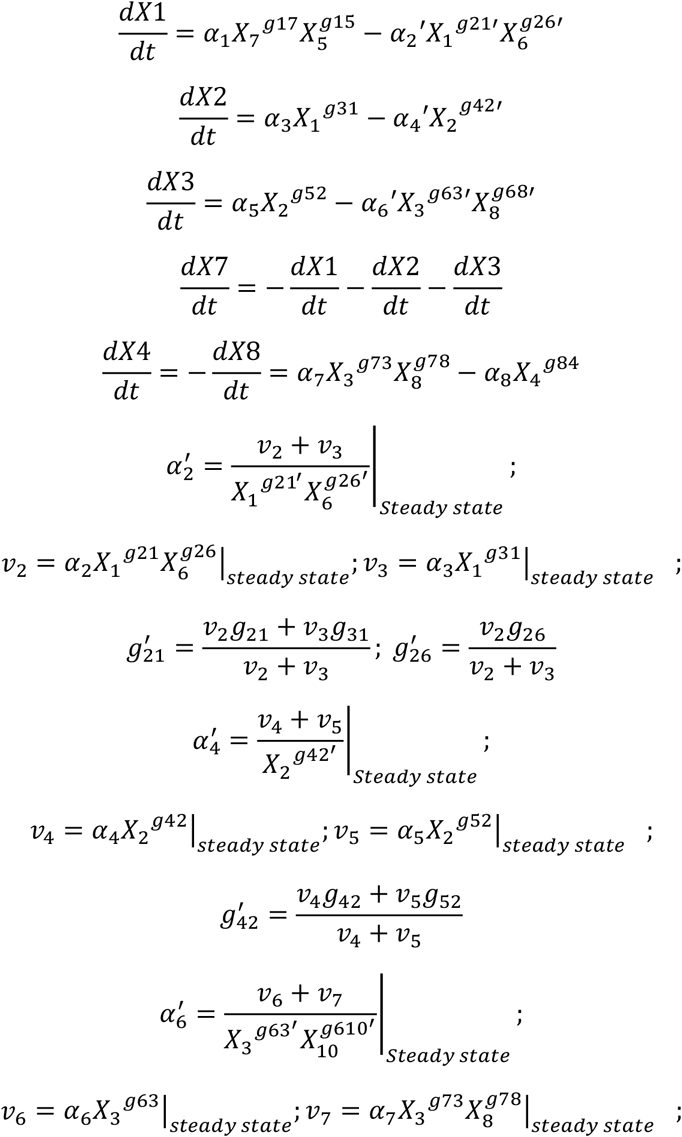

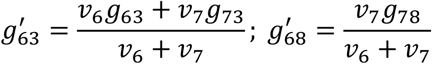

Architecture M4

The conceptual scheme we used to derive the power law mathematical model for architecture M4 is as follows

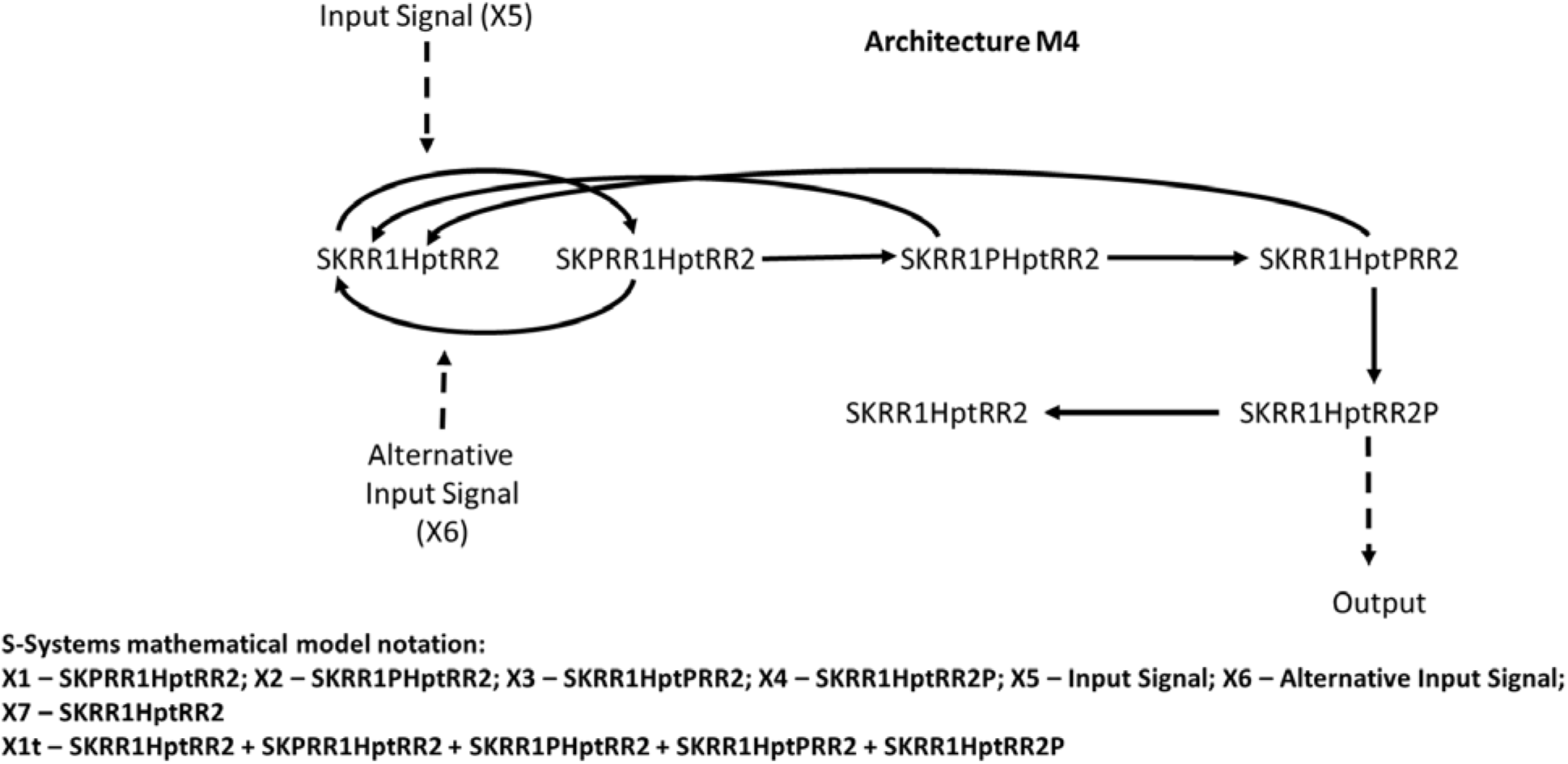

GMA Model

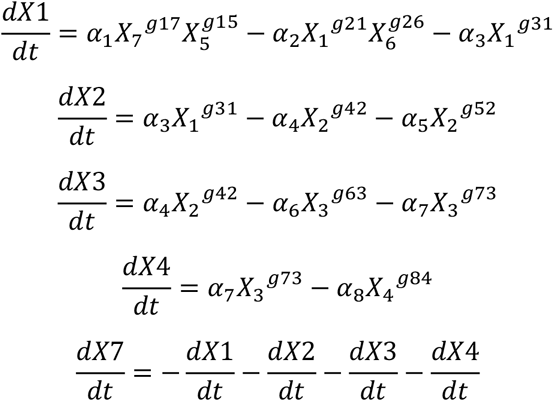

S-system Model

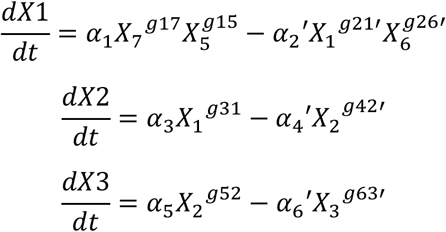

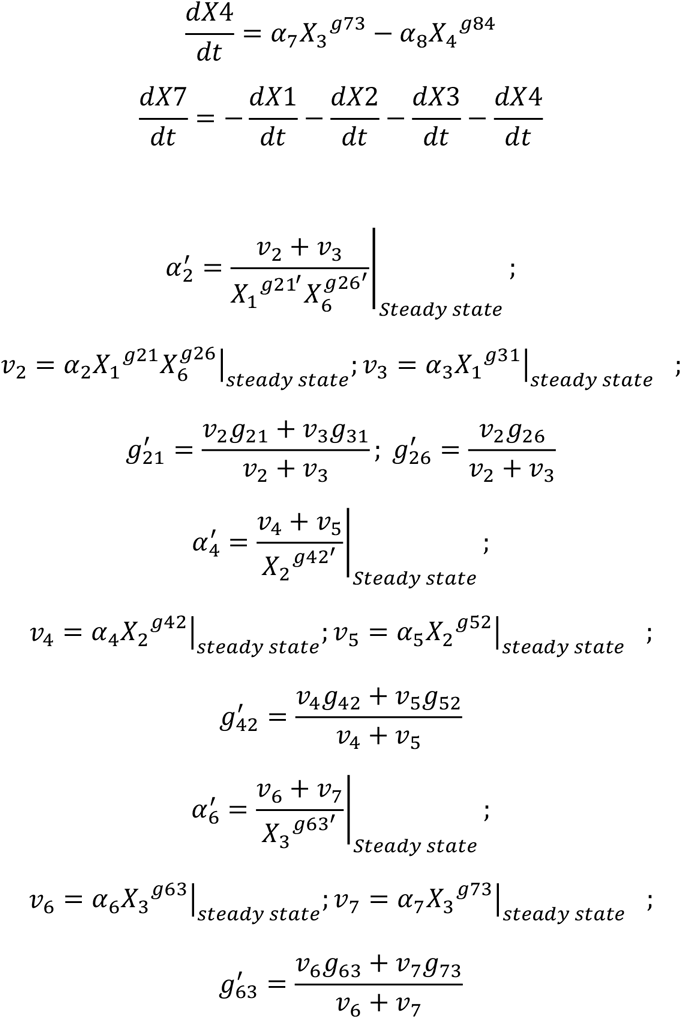

### Conceptual schemes and mass action mathematical models

#### Architecture M1

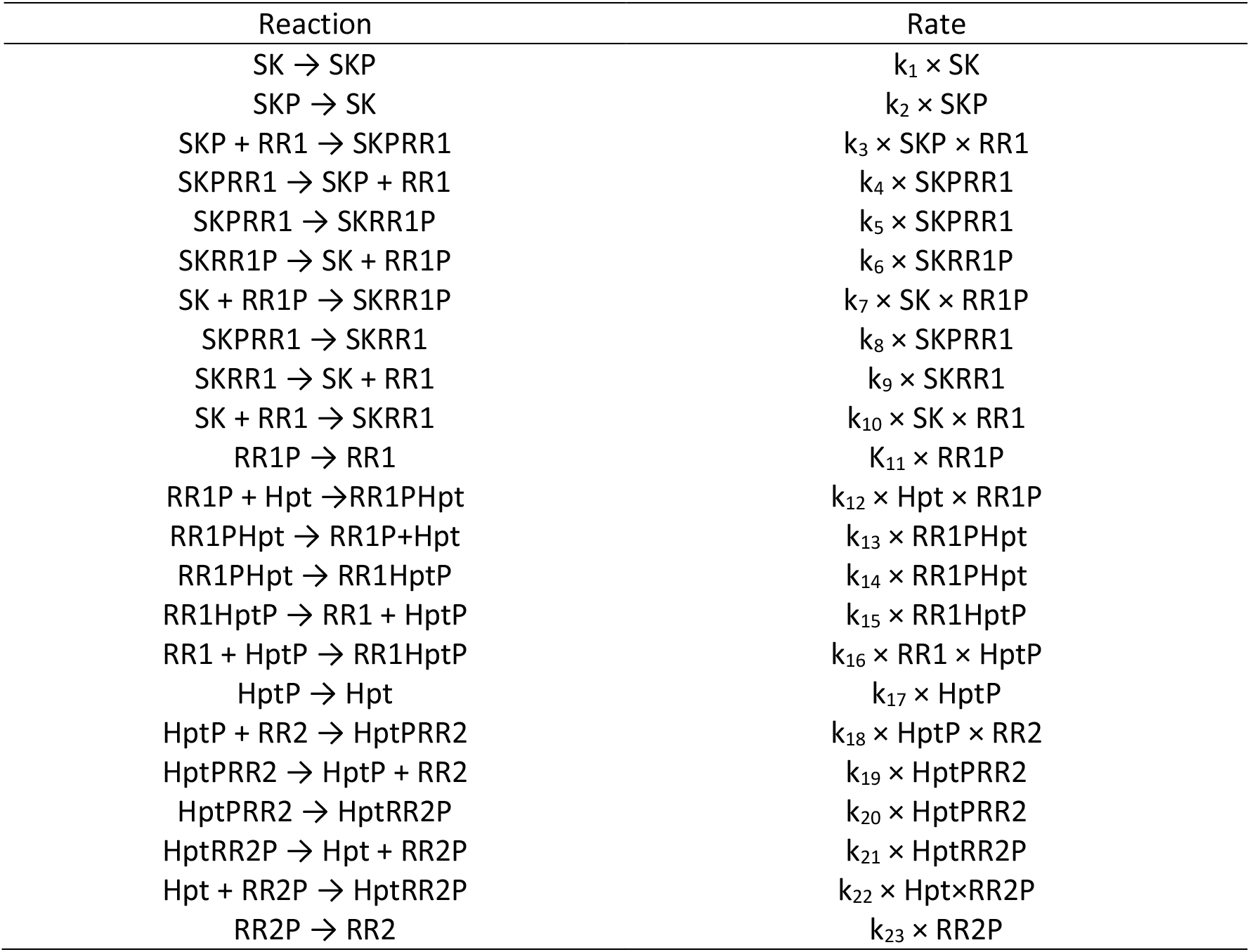

#### Architecture M2

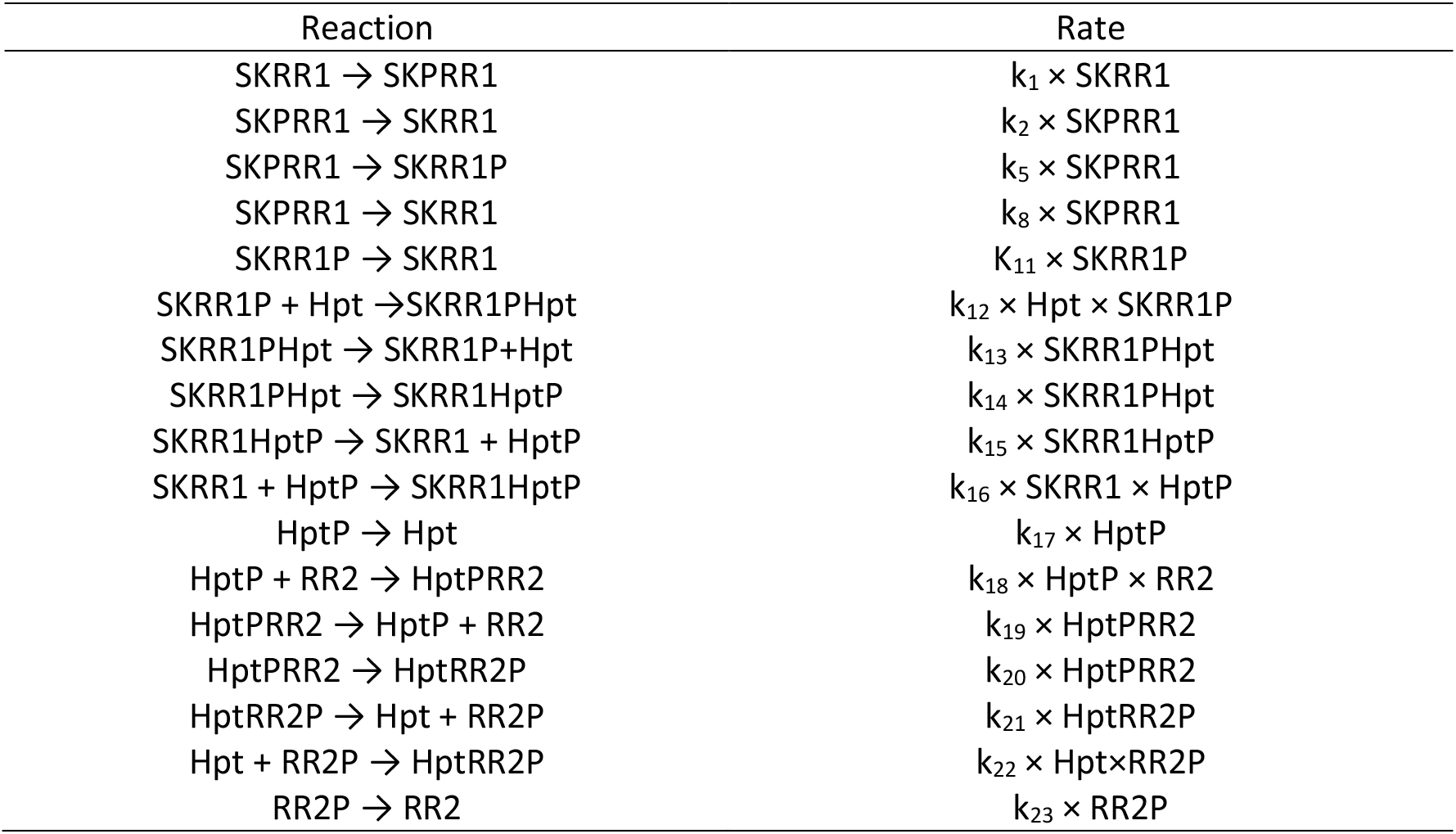

#### Architecture M2’

Architecture M2'
| Reaction | Rate |
| --- | --- |
| $SKHpt \rightarrow SKPHpt$ | $k_1 \times SKHpt$ |
| $SKPHpt \rightarrow SKHpt$ | $k_2 \times SKPHpt$ |
| $SKPHpt + RR1 \rightarrow SKPHptRR1$ | $k_3 \times SKPHpt \times RR1$ |
| $SKPHptRR1 \rightarrow SKPHpt + RR1$ | $k_4 \times SKPHptRR1$ |
| $SKPHptRR1 \rightarrow SKHptRR1P$ | $k_5 \times SKPHptRR1$ |
| $SKHptRR1P \rightarrow SKHpt + RR1P$ | $k_6 \times SKHptRR1P$ |
| $SKHpt + RR1P \rightarrow SKHptRR1P$ | $k_7 \times SKHpt \times RR1P$ |
| $SKPHptRR1 \rightarrow SKHptRR1$ | $k_8 \times SKPHptRR1$ |
| $SKHptRR1 \rightarrow SKHpt + RR1$ | $k_9 \times SKHptRR1$ |
| $SKHpt + RR1 \rightarrow SKHptRR1$ | $k_{10} \times SKHpt \times RR1$ |
| $RR1P \rightarrow RR1$ | $K_{11} \times RR1P$ |
| $RR1P + SKHpt \rightarrow RR1PSKHpt$ | $k_{12} \times SKHpt \times RR1P$ |
| $RR1PSKHpt \rightarrow RR1P + SKHpt$ | $k_{13} \times RR1PSKHpt$ |
| $RR1PSKHpt \rightarrow RR1SKHptP$ | $k_{14} \times RR1PSKHpt$ |
| $RR1SKHptP \rightarrow RR1 + SKHptP$ | $k_{15} \times RR1SKHptP$ |
| $RR1 + SKHptP \rightarrow RR1SKHptP$ | $k_{16} \times RR1 \times SKHptP$ |
| $SKHptP \rightarrow SKHpt$ | $k_{17} \times SKHptP$ |
| $SKHptP + RR2 \rightarrow SKHptPRR2$ | $k_{18} \times SKHptP \times RR2$ |
| $SKHptPRR2 \rightarrow SKHptP + RR2$ | $k_{19} \times SKHptPRR2$ |
| $SKHptPRR2 \rightarrow SKHptRR2P$ | $k_{20} \times SKHptPRR2$ |
| $SKHptRR2P \rightarrow SKHpt + RR2P$ | $k_{21} \times SKHptRR2P$ |
| $SKHpt + RR2P \rightarrow SKHptRR2P$ | $k_{22} \times SKHpt \times RR2P$ |
| $RR2P \rightarrow RR2$ | $k_{23} \times RR2P$ |

#### Architecture M3

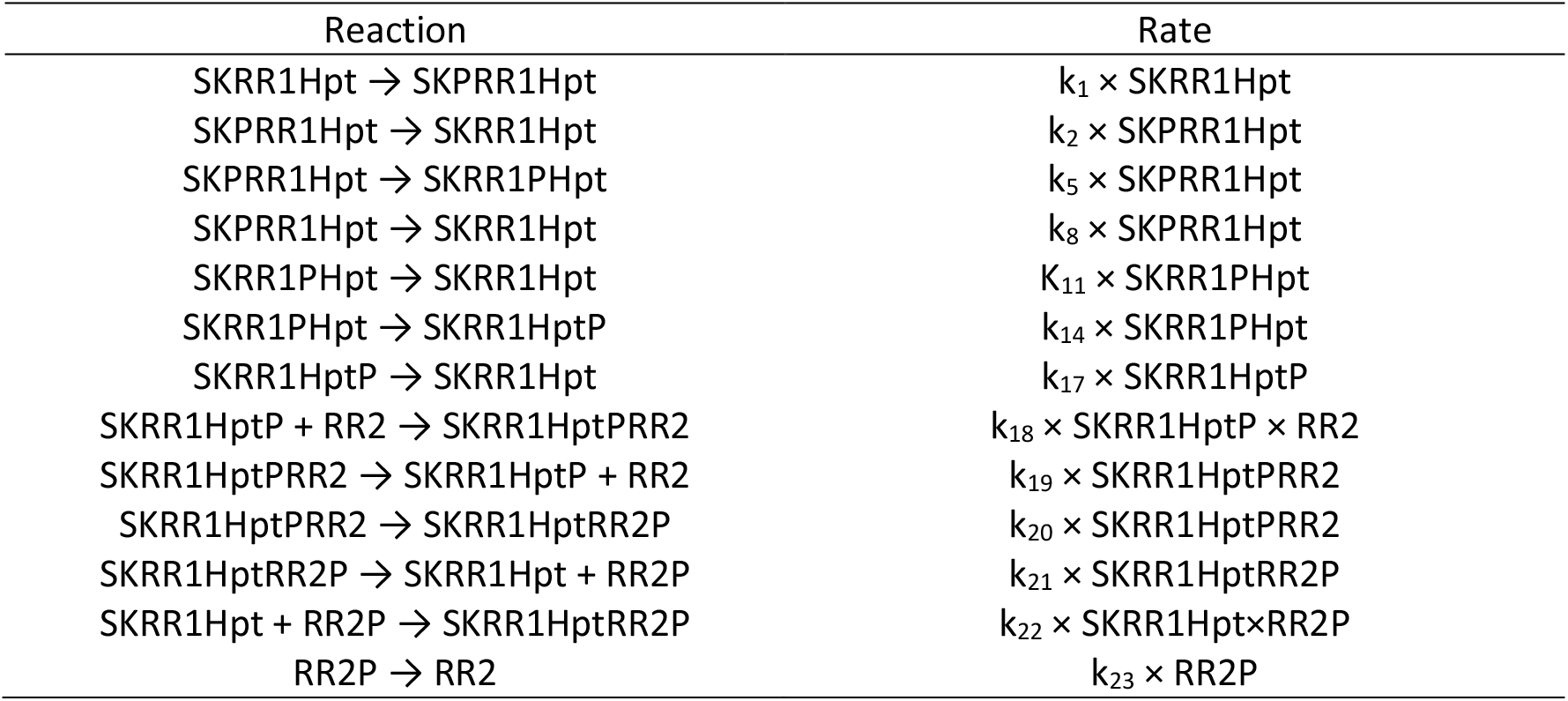

#### Architecture M4

| Reaction | Rate |
| --- | --- |
| SKRR1HptRR2 → SKPRR1HptRR2 | $k_1 \times \text{SKRR1HptRR2}$ |
| SKPRR1HptRR2 → SKRR1HptRR2 | $k_2 \times \text{SKPRR1HptRR2}$ |
| SKPRR1HptRR2 → SKRR1PHptRR2 | $k_5 \times \text{SKPRR1HptRR2}$ |
| SKPRR1HptRR2 → SKRR1HptRR2 | $k_8 \times \text{SKPRR1HptRR2}$ |
| SKRR1PHptRR2 → SKRR1HptRR2 | $K_{11} \times \text{SKRR1PHptRR2}$ |
| SKRR1PHptRR2 → SKRR1HptPRR2 | $k_{14} \times \text{SKRR1PHptRR2}$ |
| SKRR1HptPRR2 → SKRR1HptRR2 | $k_{17} \times \text{SKRR1HptPRR2}$ |
| SKRR1HptPRR2 → SKRR1HptRR2P | $k_{20} \times \text{SKRR1HptPRR2}$ |
| SKRR1HptRR2P → SKRR1HptRR2 | $k_{23} \times \text{SKRR1HptRR2P}$ |

### Parameter values and protein abundances for the mass action mathematical models

The parameter values for the mass action models were generated by combining all possible values described in Table S4. The protein abundances were approximated to the orders of magnitudes experimentally determined in Table S3 and then also combined using a latin hypercube approach.

### Mathematical Models for the Spo0 Phosphorelay

#### Parameters and abundances for the Spo0 system in *Bacillus subtilis* compiled from ^56,99^

| Reaction | Rate constants |
| --- | --- |
| SK → SKP | 0.0019 s<sup>-1</sup> |
| SKP → SK | 0 |
| SK+RR → SKRR | 100 |
| SKRR → SK + RR | 200 |
| SKP+RR → SKPRR | 200 |
| SKPRR → SKP + RR | 100 |
| SKPRR → SKRR | 0 |
| SKPRR → SKRRP | 0.08 s<sup>-1</sup> |
| SKRRP → SKRR | 0 |
| SKRRP → SK + RRP | 100 |
| SK + RRP → SKRRP | 200 |
| RRP → RR | 0.05 |
| RRP + Hpt → RRP Hpt | 100 |
| RRP Hpt → RRP + Hpt | 2300 |
| RRP Hpt → RRHptP | 1.5*10<sup>-5</sup> s<sup>-1</sup> |
| RRHptP → RR + HptP | 2300 |
| RR + HptP → RRHptP | 100 |
| HptP + RR2 → HptPRR2 | 400 |
| HptPRR2 → HptP + RR2 | 100 |
| HptPRR2 → HptRR2P | 0.20 s<sup>-1</sup> |
| HptRR2P → Hpt + RR2P | 100 |
| Hpt + RR2P → HptRR2P | 400 |
| RR2P → RR2 | 0.0001-0.006 |

| PR proteins | Abundances |
| --- | --- |
| KinA | 610 molecules cell<sup>-1</sup> |
| Spo0F | 123 molecules cell<sup>-1</sup> |
| Spo0A | 192 molecules cell<sup>-1</sup> |
| Spo0B | 267 molecules cell<sup>-1</sup> |

### Mathematical Models for the Sln1 Phosphorelay

#### Parameter values and abundances for the Sln1-Ypd1-Ssk1-Skn7 system in *Sacharomyces cerevisiae* compiled from ^30,31^

| Reaction | Sln1:Ypd1:Ssk1 | Sln1:Ypd1:Skn7 |
| --- | --- | --- |
| SKRR → SKPRR | 10000 | 10000 |
| SKPRR → SKRR | 100 | 100 |
| SKPRR → SKRRP | 10000 | 10000 |
| SKRRP → SKRR | 0.001<br>(0.0005 w/osmolites) | 0.001 |
| SKRRP + Hpt → SKRRPHpt | 2000 M<sup>-1</sup> s<sup>-1</sup> | 2000 M<sup>-1</sup> s<sup>-1</sup> |
| SKRRPHpt → SKRRP + Hpt | 2800 s<sup>-1</sup> | 2800 s<sup>-1</sup> |
| SKRRPHpt → SKRRHptP | 29 s<sup>-1</sup> | 29 s<sup>-1</sup> |
| SKRRHptP → SKRRPHpt | 7.5 s<sup>-1</sup> | 7.5 s<sup>-1</sup> |
| SKRRHptP → SKRR + HptP | 14000 | 14000 |
| SKRR + HptP → RRHptP | 2000 | 2000 |
| HptP + RR2 → HptPRR2 | 2000 | 2000 |
| HptPRR2 → HptP + RR2 | 4800 | 2800 |
| HptPRR2 → HptRR2P | 160 s<sup>-1</sup> | 1.4 s<sup>-1</sup> |
| HptRR2P → HptPRR2 | 0 | 0.4 s<sup>-1</sup> |
| HptRR2P → Hpt + RR2P | 0 | 10000 |
| Hpt + RR2P → HptRR2P | 0 | 2 |

| PR Proteins | Protein abundances |
| --- | --- |
| Sln1 | 696 molecules cell<sup>-1</sup> |
| Ypd1 | 6560 molecules cell<sup>-1</sup> |
| Ssk1 | 1018 molecules cell<sup>-1</sup> |
| Skn7 | 2536 molecules cell<sup>-1</sup> |

